# Dissociation between hemodynamic and neurochemical responses during chemogenetic modulation of cortical circuits in rats

**DOI:** 10.64898/2026.06.22.733828

**Authors:** Fatemeh Anvari-Vind, Nathalie Just

## Abstract

**Introduction:** Chemogenetic tools such as Designer Receptors Exclusively Activated by Designer Drugs (DREADDs) provide a powerful means to causally manipulate defined neuronal populations in vivo. While chemogenetic fMRI studies have consistently demonstrated robust hemodynamic responses following circuit perturbation, considerably less is known about the accompanying metabolic consequences. Functional magnetic resonance spectroscopy (fMRS) offers the potential to probe these neurochemical processes, yet the relationship between hemodynamic and metabolic responses remains poorly understood. Here, we combined chemogenetics, pharmacological fMRI (ph-fMRI), and proton magnetic resonance spectroscopy (^1^H-MRS/fMRS) at 7 T to investigate the temporal evolution of metabolic and hemodynamic responses in the rat motor cortex.

**Methods:** Female Fischer rats received viral injections in the motor cortex to express either a pan-neuronal hM3D(Gq) DREADD construct (hSyn-hM3Dq) or an interneuron-targeted construct (hDlx-hM3Dq). Ph-fMRI, fMRS, and ^1^H-MRS measurements were performed before, during, and following systemic administration of clozapine-N-oxide (CNO, 1 mg/kg). Functional MRS was acquired during the acute response phase (0–60 min post-injection), while conventional ^1^H-MRS measurements were obtained at a delayed time point (70 min post-injection).

**Results:** Chemogenetic modulation produced robust and opposing hemodynamic responses. Pan-neuronal activation elicited focal positive BOLD responses (+3.5 ± 1.5%), whereas interneuron-targeted activation generated significant negative BOLD responses (−3.3 ± 0.8%). In contrast, acute fMRS measurements revealed no significant changes in Glx or GABA concentrations during the first hour following CNO administration, despite the presence of strong hemodynamic effects. However, delayed metabolic alterations were detected 70 min after CNO administration. Animals expressing the pan-neuronal construct exhibited significant increases in GABA (+14.4%) and total choline compounds (+57.8%), whereas interneuron-targeted animals displayed reductions in several metabolites, including Glx (−15.6%), total NAA (−16.9%), glucose (−25.9%), and total creatine (−25.4%).

**Conclusion:** Chemogenetic perturbation of cortical circuits produced robust hemodynamic responses but more subtle and temporally complex metabolic effects. The absence of detectable acute changes in Glx and GABA despite strong BOLD responses, together with the emergence of delayed neurochemical alterations, highlights the challenges of interpreting metabolic signals in relation to circuit activity.

## 1. INTRODUCTION

Chemogenetic tools such as Designer Receptors Exclusively Activated by Designer Drugs (DREADDs) have revolutionized neuroscience by enabling cell-type–specific, reversible and minimally invasive modulation of neuronal activity. With DREADDs, genetically defined excitatory or inhibitory neuronal populations can be activated or silenced through systemic administration of an otherwise inert ligand (e.g., clozapine-N-oxide, CNO), allowing causal interrogation of brain circuits and behaviour without the need for chronic implants, optical fibers, or electrodes [1–2].

In parallel, Blood Oxygen Level dependent (BOLD) functional magnetic resonance imaging (fMRI) has emerged as a useful tool to monitor hemodynamic consequences of such manipulations across distributed networks. Many preclinical chemogenetic studies, especially in rodents, have employed 7-Tesla (7LT) MRI, leveraging established DREADD-fMRI protocols [3–4]. For instance, in the seminal “chemo-fMRI” study activating serotonergic neurons, region-specific BOLD responses were reliably detected following DREADD stimulation [1]. Similarly, inhibitory DREADD manipulations (e.g., targeting dopaminergic neurons) have been shown to modulate BOLD responses to external stimulation, confirming that hemodynamic readouts track network-level effects of chemogenetic perturbations [5].

These achievements have encouraged the view that DREADD-fMRI at 7LT is a robust platform for mapping functional connectivity and circuit modulation in vivo. However, BOLD is only an indirect marker of neuronal activity [6–7]. It reflects neurovascular coupling and hemodynamic changes, not the fundamental neurochemical alterations (e.g., in excitatory/inhibitory neurotransmitter pools) that drive synaptic transmission and metabolic demand. Thus, without complementary measures, BOLD fMRI alone cannot reveal whether activation or inhibition via DREADDs actually alters the balance of glutamate, γ-amino butyric acid (GABA), or other metabolic substrates in a region.

Proton magnetic resonance spectroscopy (¹H-MRS), in contrast, offers a non-invasive means to quantify neurometabolites such as glutamate (Glu) and GABA in vivo. Under appropriate acquisition parameters, ^1^H-MRS can provide estimates of regional concentrations of these key neurotransmitters/neurochemicals, arguably offering a more direct window into excitatory/inhibitory balance than hemodynamics alone. Functional MRS (^1^H-fMRS), which tracks temporal changes in metabolite concentrations during stimulation or manipulation, holds promise to reveal metabolic correlates of neural activation [8–9]. Recent human and animal fMRS studies have indeed reported small but significant changes in Glu or Glx (GluL+LGln) during sensory or cognitive tasks, suggesting that neurotransmitter pools can shift measurably under physiological stimulation [10, 11, 12]. Moreover, one elegant study combining fMRS with two-photon calcium imaging in awake mice found that MRS-derived Glu and GABA changes tracked documented shifts in excitatory and inhibitory neuronal activity, respectively bolstering the idea that MRS can reflect functional neurotransmission, not just baseline metabolic pools [13].

Despite this promise, relatively few studies have combined chemogenetics (DREADDs) with functional magnetic resonance spectroscopy (fMRS), particularly in preclinical models. While chemogenetic-fMRI studies have consistently demonstrated robust hemodynamic responses following selective circuit perturbation [3,4], considerably less is known about the accompanying metabolic consequences. Existing chemo-fMRS investigations remain scarce, and the relationship between neuronal manipulation, metabolic responses, and hemodynamic changes remains poorly understood. Moreover, recent meta-analyses of ^1^H-fMRS studies have highlighted substantial variability across paradigms, generally modest effect sizes for Glu/Glx, and often inconsistent GABA responses, emphasizing the influence of stimulus design, brain region, acquisition strategy, and temporal resolution on metabolite detectability [14].

Importantly, these observations raise a broader question that extends beyond methodological sensitivity: what does a metabolic signal actually reveal about neural circuit function? While BOLD fMRI provides an indirect measure of activity through vascular responses, fMRS offers the possibility of probing the neurochemical processes that support neuronal signaling. However, the precise relationship between hemodynamic and metabolic responses remains incompletely understood. Detectable changes in Glu, Gln, GABA, or related metabolites may reflect alterations in neurotransmitter cycling, cellular energetics, glial-neuronal interactions, or network reorganization, but their interpretation in terms of circuit activity is still debated. Chemogenetic approaches provide a unique opportunity to address this question. By selectively modulating defined neuronal populations, DREADDs allow causal perturbation of neural circuits while simultaneously monitoring their hemodynamic and metabolic consequences. Combining chemogenetics with fMRI and fMRS therefore offers a powerful framework for investigating how circuit-level manipulations are translated into vascular and metabolic responses in vivo and for probing the relationship between neurovascular and neurometabolic coupling.

In the present study, we applied this multimodal strategy to the rat motor cortex, a region extensively investigated using chemogenetic fMRI at 7 T. Two complementary chemogenetic approaches were employed. First, a pan-neuronal hSyn-hM3D(Gq) construct was used to activate the local neuronal population broadly. Second, an hDlx-hM3D(Gq) construct was used to selectively activate GABAergic interneurons, thereby increasing inhibitory tone within the same cortical network. These complementary perturbations enabled us to investigate how distinct cellular drivers of circuit activity are reflected in hemodynamic and metabolic measurements. Using concurrent assessment of BOLD responses, interhemispheric communication and metabolic dynamics with ^1^H-fMRS, we sought to determine how metabolic signals relate to causal manipulations of cortical circuitry. More broadly, we asked whether hemodynamic and metabolic measurements provide complementary or divergent information regarding neural circuit function. Finally, computational simulations at higher magnetic field strengths were performed to explore the detectability of plausible neurometabolic changes under improved spectral resolution and signal-to-noise conditions. Together, these experiments provide a framework for interpreting metabolic measurements in the context of circuit-level perturbations and highlight both the opportunities and challenges of combining chemogenetics, fMRI and fMRS.

## 2. MATERIALS and METHODS

### 2.1. Animals

Female Fisher rats (Janvier, France; BW= 190±20g) were used in the present study. Rats were housed in pairs in the Danish Research Centre for Magnetic resonance (DRCMR) animal facility with controlled temperature (18–22°C) and humidity (50–56%) and free access to food and water under a 12-h reverse light-dark cycle (light off at 19:00). Two weeks after their arrival within the facility, rats were transported to the genetically modified (GMO) laboratory for stereotactic viral injections.

For general activation of cortical neurons within motor cortical areas (M_1,2_), we used the pAAV8-hSyn-hM3D(Gq)-mCherry construct. The pAAV8-hSyn-hM3D(Gq)-mCherry vector enables robust pan-neuronal expression of hM3D(Gq), supporting effective DREADD activation. While the hSyn promoter drives expression specifically in neurons, it does not distinguish between excitatory and inhibitory subtypes, potentially complicating interpretations in heterogeneous cortical regions and this limits our ability to dissect cell-type-specific contributions. Nonetheless, this broad activation was appropriate for our primary objective of validating chemo-fMRS as a tool for studying cortical neurochemical dynamics. Selective promoters may be used in future studies to dissect cell-type-specific contributions. To induce inhibitory modulation of cortical circuits, we used the pAAV-hDlx-Gq-DREADD-dTomato-Fishell-4 construct, which drives expression of the hM3D(Gq) receptor selectively in GABAergic interneurons [15]. Upon CNO administration, activation of these interneurons is expected to increase inhibitory tone and thereby reduce the activity of neighboring excitatory neurons within the local cortical network. These AAV were produced with a retrograde serotype, which permits retrograde access to projection neurons. For the sham group, animals underwent identical stereotaxic procedures and received injections of the AAV5-CaMKIIa-hChR2(H134R)-EYFP construct. This construct was selected as a viral expression control because it results in stable neuronal transgene expression and fluorescent labeling of transduced cells without chemogenetic activation in the absence of optical stimulation. Consequently, these animals controlled for potential effects related to surgery, viral transduction, transgene expression, and fluorescent protein expression while lacking the capacity for CNO-induced neuronal modulation. In the absence of optical stimulation, ChR2 expression does not induce neuronal activation and therefore served as a control for surgical procedures, viral transduction, and long-term expression of an exogenous membrane protein. This control was not intended to mimic DREADD signaling but rather to account for non-specific effects associated with viral delivery and transgene expression. Control rats (CTL) with or without viral transductions were also used in conjunction with saline (0.9%) or Clozapine-N-Oxide (CNO) to serve as a reference for physiological, pharmacological and neurochemical effects. For surgeries, rats were mounted in a stereotaxic frame under isoflurane anesthetics. Default isoflurane concentration for the isoflurane-oxygen mixture used was 2%, with the concentration being adjusted during the surgery. 10 minutes prior to the surgical procedure a 0.15 ml lidocaine/bupivacaine solution (lidocaine 10 mg/ml, 10%; bupivacaine 5 mg/ml, 20%) was applied locally (in sterile water), combined with 0.15 ml systemic injections of norodyl (50 mg/ml, 10% dilution), baytril (50 mg/ml, 10% dilution) and buprenorphrine (0.3 mg/ml, 10% dilution) solutions (in saline). Stereotaxic coordinates were chosen to target the injections at a specific confined portion of the motor cortex (M_1,2_) (AP: 0-1 mm from bregma; ML: 2.0-3.2 mm from bregma). A small craniotomy was made on the right hemisphere; the syringe was slowly inserted 1 mm below the cortical surface and 1μl of the viral construct was infused (0.2μl/min). Following surgeries, animals received postoperative treatment and were housed for a minimum of 4 weeks, to allow for sufficient expression of the genes encoding for the receptors. A total of 60 Fisher rats were used as described in Table 1.

**Table1:** Number of Animals in different groups and for histology/Immunohistochemistry.

| **Group** | **CTL (+Saline)** | **Sham** | **pAAV8-hsyn-**<br>**hM3D(Gq)_mcherry**<br>**(eDREADDs)** | **pAAV-hdlx-Gq-**<br>**DREADD-**<br>**dTomato_Fishell4**<br>**(iDREADDs)** | **Histology/IHC** |
| --- | --- | --- | --- | --- | --- |
|  | 6 | 7 | 8 | 10 | 29 |

### 2.2 MRI acquisition

All *in-vivo* MRI experiments were carried out in a 7T Bruker-Biopsec 70/20 animal scanner (Bruker, Wissembourg, France) using an 86 mm diameter volume coil for transmission (Bruker, Ettlingen, Germany) and a single loop 20-mm surface coil for reception (Bruker, Ettlingen, Germany). Unconsciousness was induced in a chamber filled with 5% isoflurane and maintained throughout preparation with 2-3% isoflurane in air and oxygen (70/30%) supplied through a nose cone. A cannula was inserted in the left quadrant of the rat’s intraperitoneal area for subsequent infusions of 1mg/kg Clozapine-N-oxide (CNO). The animal was placed in a supine position with its head secured in a stereotactic frame equipped with a bite bar and ear bars. The eyes were covered with ophthalmic ointment to prevent them from drying. Physiological monitoring equipment consisted of a rectal probe for continuous temperature recording (37± 0.5°C), a pneumatic sensor placed underneath the rat’s body (SA Instruments, NY, USA). Further, both the partial pressure of O_2_ (SpO_2_) and heart rate was continuously monitored through an oximeter attached to the left hind paw (Nonin Medical B.V, The Netherlands). Continuous readings were obtained through in-house written Python scripts. Once the respiratory rate and temperature of the rat were stabilized, a bolus of medetomidine (0.05 mg/kg; DexDomitor, Orion corporation, Finland) was delivered through a subcutaneous syringe attached to the left flank of the rat. Following a decrease of the respiratory rate and of the heart rate, isoflurane anesthesia was reduced to 0.25% and 10 minutes after the end of the bolus injection, medetomidine infusion (0.1mg/kg/h) was started based on the respiratory rate of the rat. Physiological parameters were monitored throughout all MR experiments. Blood oxygen saturation (SpOL) was maintained between 92% and 100% across all animals. Bolus administration of medetomidine led to an initial HR reduction in most rats, with subsequent recovery profiles varying across individuals either sharp, gradual, or absent. These trends were observed in DREADDs, sham, and control (CTL) groups. Notably, only DREADD animals exhibited additional HR fluctuations following CNO injection, suggesting an effect of chemogenetic activation on autonomic regulation. Since we are also interested in the characterization of interhemispheric mechanisms, we decided to proceed with experiments without intubation [16].

The outline of experiments is described in Figure 1. First, structural T_2_-weighted TurboRARE images covering the rat brain were acquired (TR/TE=2700/33 ms; RARE factor = 8; Matrix=256 x 256; FOV= 25 x 25 mm; number of slices = 25; Slice Thickness = 0.8 mm). One hour after medetomidine induction, pharmacological fMRI (Ph-fMRI) scans were acquired using a gradient-echo echoplanar imaging (GRE-EPI) sequence aligned to the previously acquired structural images (TR/TE/flip angle=1000/11 ms/60°; Matrix=64 x 64; FOV= 25 x 25 mm; repetitions: 315; Slice thickness = 0.8 mm). Pharmacological fMRI (Ph-fMRI) was conducted with 3000 repetitions (total acquisition time = 59 minutes). The bolus of CNO was performed manually 15 minutes after the start of the EPI acquisition.

**Figure 1.**
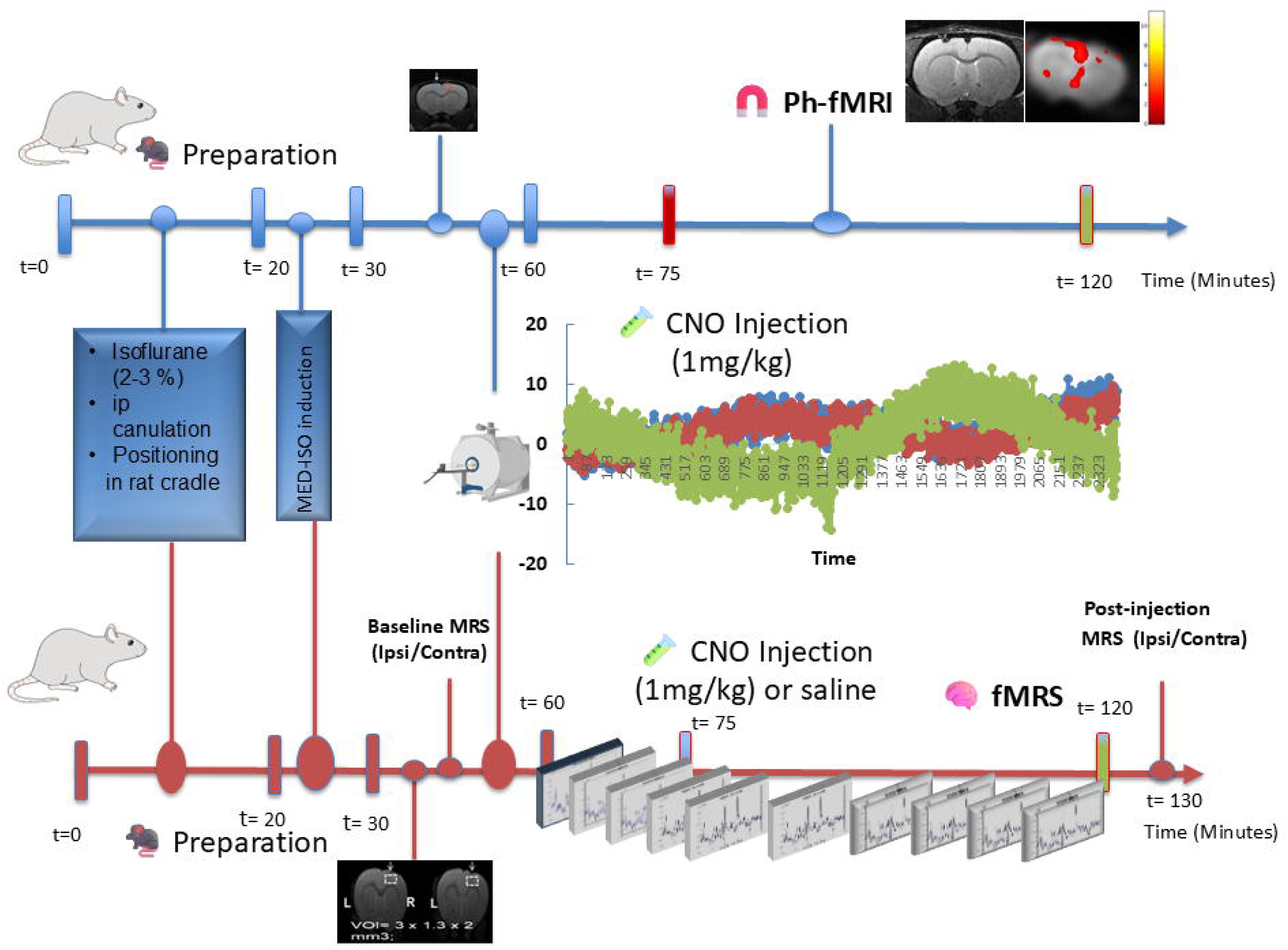
Timelines summarizing anesthesia (ISO→MED-ISO), baseline, Ph-fMRI. (1) with CNO 1 mg/kg at t=15 min, dynamic ¹H-fMRS (0–60 min, 2), and delayed static ¹H-MRS (∼70 min post-CNO) acquired ipsilateral and contralateral to the injection site.

### 2.3 Image processing

#### 2.3.1 Preprocessing of fMRI series

All MR images were first exported from ParaVision 6.0.1 to DICOM format and then converted to NIfTI (Neuroimaging Informatics Technology Initiative; http://nifti.nimh.nih.gov) using the dcm2nii tool (https://www.nitrc.org/projects/dcm2nii/). The structural image of one of the rats was chosen as the target reference space and co-registered to the SIGMA rat brain template [17]. Every other structural image was registered to the target using statistical parametric mapping (SPM12) and to the SIGMA rat template. The fMRI image series were preprocessed using functions from the SPM12 and FSL (FMRIB Software Library, https://fsl.fmrib.ox.ac.uk/fsl/fslwiki/) neuroimaging toolkits. Motion-corrected images were corrected for slice timing with SPM12 and realigned. After co-registration and normalization to the structural template spatial smoothing was performed using a 1Lmm 3D Gaussian kernel using SPM12 (MATLAB; The Mathworks, Natick, MA, USA; Statistical Parametric Mapping, www.fil.ion.ucl.ac.uk/spm/).

#### 2.3.2 Mapping BOLD responses

For Ph-fMRI processing, in-house Matlab routines were used. A ROI-based analysis was conducted to retrieve the dynamic GRE-EPI signal intensity time courses. ROIs were drawn over the chemo-induced area within the MC (M_1,2_) of each rat brain and its symmetrical homotopic area. Other ROIs were drawn in the somatosensory cortices and in subcortical areas. In addition, we proceeded as described in [18, 19] to obtain BOLD maps during CNO challenge. Briefly, whole brain analyses were conducted to extract signal-intensity time series. We identified all the voxels that showed more than 1% change in signal intensity by performing a t-test. The average signal time course in activated voxels was determined to decide the shape of brain signal change per animal. Individual time courses were then averaged to obtain the basic shape of the to-be-modeled waveform. This waveform was used to set up the framework of the general linear model (GLM) using SPM12, with 1.2 mm FWHM spatial smoothing and a high pass filter of 3600 s. The first time series volume was deleted from further analysis to ensure steady state imaging has been reached. The design matrix consisted of the custom-made waveform explained above with an initial off period of 900 s (7 volumes of baseline recordings), followed by a gradual onset of 900 s (starting with the volume during which the CNO challenge was injected) and then a steady state on period of another 1800 s. For each subject, a custom block design matrix was constructed to model the temporal profile of the CNO-induced activation. The model consisted of three main conditions:1) Baseline (Pre-injection): A 900-second period corresponding to the initial 7 volumes before CNO administration, used to establish steady-state baseline signal. 2) Transition (Onset Phase): A 900-second period immediately following the injection, modeled as a gradual onset to account for the systemic and neural uptake delay of CNO. This was modeled using a ramp function, convolved with SPM’s canonical hemodynamic response function (HRF). 3) Activation (Steady-State CNO Effect). The remaining 1800 seconds (∼30 minutes), modeled as a plateau representing the sustained effect of CNO after reaching systemic equilibrium. Each condition was represented as a regressor of interest, and the design matrix included: the CNO challenge waveform (baseline, onset, activation), Six motion parameters (translation and rotation) as nuisance regressors, High-pass filtering (cutoff = 3600 s) to remove low-frequency drifts. Statistical inference was performed at the single-subject (first-level) using t-contrasts comparing the activation period against baseline. T statistical maps were corrected for multiple comparisons using family-wise error correction (FWE) at p < 0.05.

### 2.4 ^1^H-fMRS and static ^1^H-MRS pre and post- CNO

#### 2.4.1 Functional Proton Magnetic Resonance Spectroscopy (^1^H-fMRS)

Localized functional proton MR spectroscopy (^1^H-fMRS) was conducted using a LASER sequence (TR = 2 s; TE = 24.5 ms, Spectral width = 4000 Hz; 2048 points) modified from a sequence generously shared by Dr Julien Valette (Commissariat à l’Energie Atomique, Paris-Saclay; France). Prior to ^1^H-fMRS acquisitions, we acquired B_0_ field map and water spectrum, adjusted basic frequency, shimmed globally and locally, and optimized RF power. A large voxel of interest (VOI = 1.4 x 2.8 x 2 mm^3^) was positioned onto the anatomical T_2_-weighted images over the viral injection site covering the entire motor cortex (MC) encompassing M1 and M2 areas. After shimming using MAPSHIM and water suppression with VAPOR [20], 200 fids were acquired continuously (with 8 or 16 averages, 16 s or 32 s per time point) using the LASER sequence. After 15 minutes, CNO (1mg/kg) or saline was injected, and fids were acquired up to 59 minutes. For quantification purposes, a water spectrum was also acquired prior to each dynamic acquisition (8 fids). MAPSHIM was used for shimming over the VOI down to a linewidth of 10.7± 2 Hz.

Pre and Post- CNO ^1^H-MRS were acquired successively in the ipsilateral and contralateral homotopic MC within a voxel of identical size following re-shimming. The LASER sequence was used using similar acquisition parameters to ^1^H-fMRS and 512 averages.

#### 2.4.2 ^1^H-fMRS Data Processing

^1^H- fMRS and ^1^H- MRS data were processed using in-house written Matlab routines adapted from FID-A [21]. After Fourier transformation, individual spectra were zeroth order phase adjusted and re-aligned. After visual inspection and removal of phase outliers, spectra were first order phase adjusted and finally averaged. Metabolic time courses were obtained by averaging spectra over 15 minutes per successive time -period (baseline, transition, or activation) or 5 minutes per time point to obtain full time courses. Each point was transferred to LCModel [22] (http://www.s-provencher.com/pages/lcmodel.html). The quantification was performed with a basis-set of 20 brain metabolite spectra simulated using a density matrix procedure with FID-A [21]: Scyllo-Inositol (Scyllo); Ascorbate (Asc); Alanine (Ala); Aspartate (Asp); Glycerophosphocholine (GPC); Phosphocholine (PCh); Creatine (Cr); Phosphocreatine (PCr); γ-amino-butyric acid (GABA) ; Glucose (Glc); Glutamine (Gln);Glutamate (Glu); myo-Inositol (Ins); Lactate (Lac); N-Acetyl-Aspartate (NAA); N-acetylaspartyl-glutamic acid (NAAG); Phosphatidylethanolamine (PE); Taurine (Tau); total creatine (tCr: PCr + Cr, total NAA (tNAA= NAA + NAAG) and Glx = Glu + Gln were also assessed.

A spectrum of fast-relaxing macromolecules was also included in the basis set: Macromolecules (Mac). This macromolecule spectrum was measured over a 4 x 4 x 4 mm^3^ brain VOI in six healthy Fisher rats with the double-inversion procedure included in the LASER sequence package with a TR= 8s. Absolute metabolite concentrations were obtained using unsuppressed water signal (eight averages) as an internal reference assuming a brain water content of 80%. The Cramer-Rao lower bounds (CRLB) were used as a reliability measure of metabolite concentration estimates. Higher concentration metabolites (Glu, NAA, Ins, Taurine, PCr and Cr) with CRLB under 5% were considered reliably quantified whereas for Glc, GABA, Lac, and Asp, CRLB under 40% were considered acceptable and data were kept for further analysis. All data including those above the defined threshold were displayed. To determine the quality of spectra, NAA and tCr (PCr + Cr) spectral linewidths and Signal to noise ratio (SNR) were also assessed. The spectral SNR was calculated as the ratio between the NAA peak amplitude and the standard deviation of spectral noise measured between 5 and 8 ppm.

#### 2.4.3 Metabolite Concentration time courses

In-house written Matlab routines were used. Individual fids were Fourier transformed. 20 individual spectra, representing 5 minutes of acquisition were realigned, phased, and averaged allowing us to obtain 10 successive time points per metabolite. 5-min averaged spectra were transferred to LCModel and adjusted for each animal. Mean metabolite time courses were obtained by averaging metabolite concentrations across animals for eDREADDs, iDREADDs, Sham and controls. The spectral SNR was calculated as the ratio between the NAA peak amplitude and the standard deviation of spectral noise measured between 5 and 8 ppm.

In order to explore further the impact of DREADD activation or inhibition in both hemispheres of the brain, successive ^1^H -MR Spectra were further acquired in the same chemo-induced ipsilateral VOI used for chemo-fMRS and in its homotopic contralateral part, prior to CNO injection and 70-min post-CNO injection. This timepoint was chosen because systemic and brain concentrations of CNO become measurable within 15–30 minutes after systemic administration and downstream neurometabolic effects detectable by MRS typically evolve over tens of minutes; performing MRS at ∼70 minutes therefore balances early receptor engagement with the time required for metabolic shifts to become measurable while minimizing later systemic/off-target effects (for example clozapine back-conversion) reported at later post-injection intervals. Ipsilateral vs contralateral comparisons were used as within-subject controls; Sham animals were included to control for systemic and off-target effects.

### 2.5 Histology and c-Fos Immunohistochemistry

To verify viral expression and anatomical specificity of the DREADD injections, dedicated animals from each group were used for histological analysis. Animals were transcardially perfused with phosphate-buffered saline (PBS, 0.01 M), followed by 4% paraformaldehyde (PFA). Brains were post-fixed in 4% PFA, cryoprotected in 30% sucrose, and sectioned coronally (40–50 µm) using a cryostat (Leica CM3050 S). Sections were mounted on gelatin-coated slides and cover slipped with ProLong Gold mounting medium. Fluorescent reporter expression was assessed using epifluorescence microscopy, and representative sections covering the full extent of the injection site were digitized using a Zeiss Axioscan Z1 slide scanner. In a subset of animals expressing excitatory DREADDs, neuronal specificity was further confirmed by immunostaining for NeuN using standard immunohistochemical procedures.

For assessment of neuronal activation, c-Fos immunohistochemistry was performed in a cohort of Fischer rats following chemogenetic stimulation. After a minimum of four weeks to allow for viral expression, animals received systemic CNO injections under light isoflurane anesthesia and were perfused approximately two hours later. Only animals receiving 1 mg/kg CNO are reported here. Free-floating coronal sections were processed using standard blocking and antibody incubation protocols, with mouse anti-c-Fos primary antibody and Alexa Fluor–conjugated secondary antibody. Sections were imaged using a Zeiss LSM 880 confocal microscope, acquiring z-stacks at 1 µm resolution.

The hDlx enhancer has been extensively validated as a tool for selective targeting of forebrain GABAergic interneurons. Previous studies demonstrated that hDlx-driven hM3D(Gq) expression is restricted to interneurons and that CNO administration selectively increases interneuron activity, resulting in enhanced inhibitory drive onto neighboring excitatory neurons [15].To provide additional confirmation of DREADD-mediated neuronal activation, c-Fos-positive cells were quantified in a limited subset of representative animals for which confocal datasets were available. Cell counts were performed bilaterally within the motor cortex and expressed as c-Fos-positive cells per mm² using ImageJ (NIH) in predefined regions of interest, including ipsilateral and contralateral motor cortex. Given the exploratory nature of this analysis and the limited sample size, results were used for qualitative comparison only

### 2.6 Biophysical simulation of metabolite dynamics at 7T and 14T (see Simulated Detectability of Chemogenetic metabolic…)

To interpret the experimental ^1^H-fMRS findings and assess field-strength–dependent detectability, we implemented a biophysical simulation framework (Python) to generate synthetic ^1^H-fMRS time courses at 7T and 14T. The model combined (i) a pharmacokinetic component describing systemic ligand dynamics and (ii) a compartmental neurometabolic system representing excitatory neurons, inhibitory neurons, and astrocytes, inspired by established glutamate–glutamine cycle and neuroenergetic models [23, 24, 25, 26]. Chemogenetic effects were implemented as fractional modulations of compartment-specific metabolite pools driven by ligand concentration through a Hill-type relationship, analogous to receptor–effector coupling models commonly used in systems pharmacology [27]. Identical underlying biological perturbations were imposed at both field strengths, while field-dependent observability was modeled by incorporating realistic differences in spectral resolution and signal-to-noise ratio. At 7T, metabolite pooling was enforced (Glx = Glu + Gln) and higher noise levels were applied, consistent with preclinical fMRS limitations [25, 28], whereas at 14T individual Glu, Gln, and GABA concentration time courses were retained with improved SNR and reduced variance [29, 30]. Inter-subject variability was simulated by randomizing key physiological parameters across virtual subjects. This framework allows direct comparison of the temporal detectability of chemogenetically induced metabolic changes at 7T versus 14T under otherwise equivalent biological conditions. Importantly, the ligand-driven component can be parameterized either to reflect acute CNO-driven activation or to reproduce delayed profiles associated with CNO–clozapine conversion, enabling interpretation of both early null effects and late metabolic changes observed experimentally.

### 2.7 Statistical Analysis

Statistical analyses were performed using R (version 4.5.2) and the lme4, lmerTest, emmeans, and effect size packages or in Prism (GraphPad, San Diego, CA). Data are presented as mean ± standard deviation (SD) unless otherwise stated. BOLD responses and static metabolite concentrations were compared across groups using a repeated measures ANOVA followed by a Bonferroni post hoc test. For each group, mean BOLD responses in contralateral and ipsilateral sides were extracted at each time point. The contra–ipsi difference was computed as diff = mean_contra - mean_ipsi.

To compare the temporal evolution of contra–ipsi differences across groups, we fitted a weighted linear model: diff“- group X time.

This model allowed testing: 1) Contra–ipsi difference within each group (intercept at time 0), 2) Differences between groups at time and 3) Group-specific temporal changes (interaction group*time). Significance was assessed at α = 0.05. Model predictions and standard errors were used for visualization of temporal patterns.

A linear mixed-effects model was fitted to assess the effects of Group (eDREADD, iDREADD, Sham, CTLs), Hemisphere (ipsilateral vs. contralateral), and Time Point on the measured metabolite concentration time courses. Fixed effects included Group, Hemisphere, TimePoint, and their interactions, while Subject was modeled as a random intercept to account for repeated measures. Model assumptions were verified via residual diagnostics. Estimated marginal means and pairwise contrasts were computed using the emmeans package with Tukey adjustment for multiple comparisons. Confidence intervals for fixed effects were obtained via profile likelihood, and standardized effect sizes (Cohen’s *d*, partial R²) were calculated using the effect size package. A p < 0.05 was considered significant.

## 3. RESULTS

### 3.1 CNO induced significant BOLD responses in the chemo-induced motor cortex

Representative c-Fos quantification performed in a subset of animals revealed increased ipsilateral neuronal activation following chemogenetic stimulation (Fig.2 A). Animals expressing the pan-neuronal hSyn-hM3D(Gq) construct exhibited the largest increase in c-Fos-positive cells within the injected motor cortex compared with the contralateral hemisphere (Fig. 2 C-F). A more moderate ipsilateral increase was observed in animals expressing the interneuron-targeted hDlx-hM3D(Gq) construct (Fig. 2H-I), whereas sham animals showed little hemispheric asymmetry and *no cfos label presence at the injection site (M1/M2) or more medial neocortical regions.* (Fig. 2G). Although based on a limited number of animals, these findings are consistent with successful chemogenetic modulation of cortical activity and support the interpretation of the imaging results.

**Figure 2.**
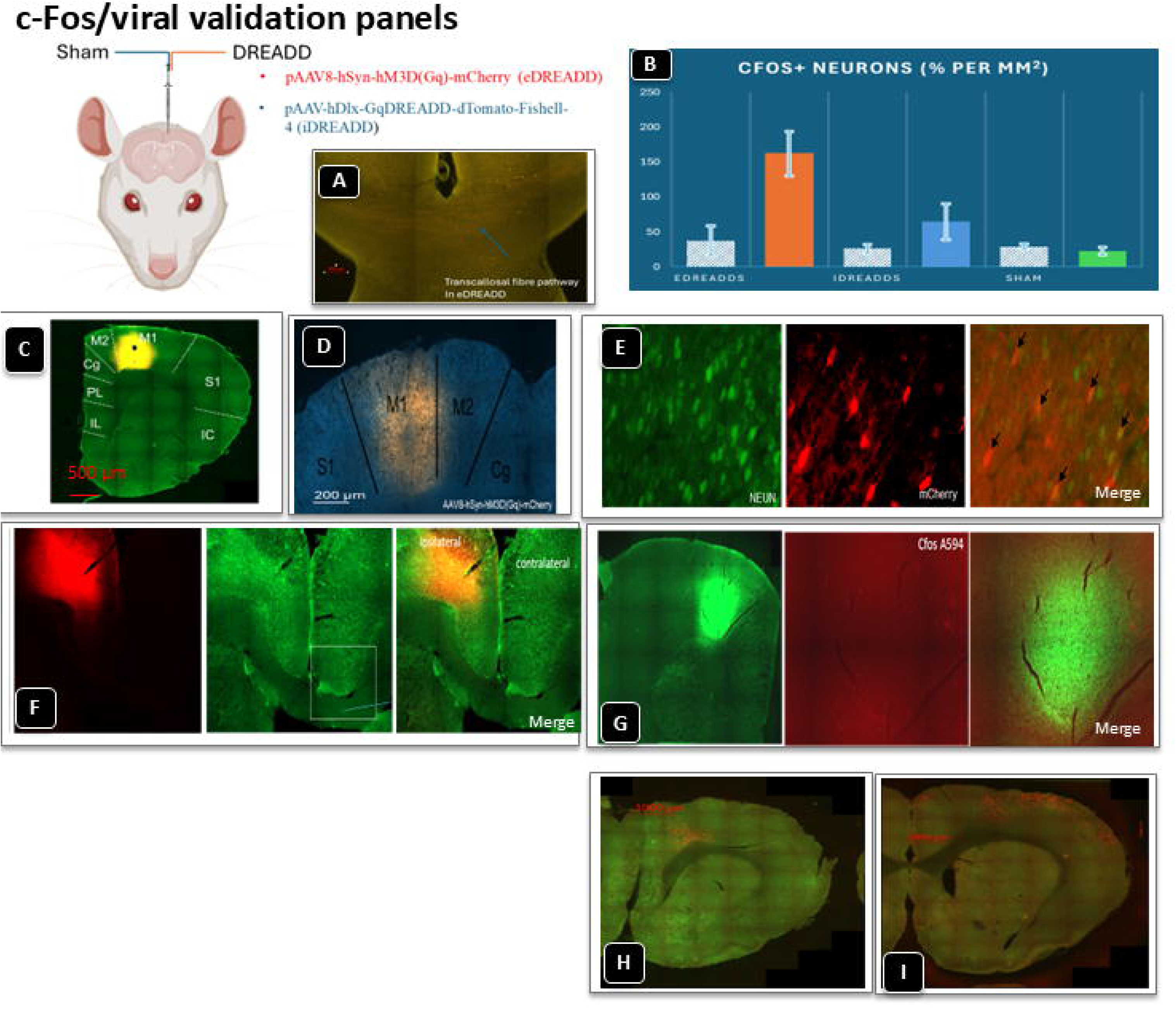
c-Fos/viral validation (M1 only). (A) Schematic/example of the transcallosal connection between homotopic MC regions for anatomical context. (B) Representative c-Fos labeling densities for a subset of rats of each group comparing contralateral versus ipsilateral sides (eDREADDs ipsilateral side : orange; iDREADDs ipsilateral side: blue; sham ipsilateral side: green). (C) Reporter fluorescence confirms viral expression restricted to M1 (AP 0–1 mm; ML 2.0–3.2 mm; depth ∼1 mm) (D) representative fluorescence images from motor cortex M1 for eDREADDs (n=12); (E) Staining for the neuronal marker NEUN and m-cherry indicated the proportion of cells infected with the AAV8-hSyn-hM3D(Gq)-mCherry virus and merging of the two (arrows indicate double-labelled cells (cells that co-express G(q)-DREADDS and NEUN)) (F) Cfos expression in contralateral and ipsilateral rat cortex, and (G) Sham animals, here provided solely to verify targeting/activation; not a primary outcome. (H-I) Representative and qualitative cFos staining for pAAV-hDlx-Gq-DREADD-dTomato-Fishell-4 construct (iDREADDs) in the rat cortical areas.

To assess how chemogenetic stimulation of the right MC via DREADDs affects regional activation, we conducted Ph-fMRI during CNO infusion after a 15-minute baseline acquisition (Fig.3). The mean ipsilateral BOLD time courses during CNO uptake in MC of eDREADDs, iDREADDs, Shams and Saline controls (CTL) demonstrated significant differences across groups (Fig.3 A). A two-way ANOVA with repeated measures was performed to compare the effects of Group (eDREADDs, iDREADDs, Shams, and CTL) and Time on the measured outcome. There was a significant Group × Time interaction (F_33,288_ = 119.17, p < 0.0001), indicating that the effect of Group differed across time points. Group alone had a significant effect (F_3,288_ = 1483.29, p < 0.0001) and Time also significantly influenced the outcome (F_11,288_ = 49.67, p < 0.0001). The significant interaction suggests that comparisons between Groups at individual time points are more informative than overall main effects. Therefore, Bonferroni corrections were applied at each time point showing a highly significant difference (p<0.001) between eDREADDs and iDREADDs time courses between 30 and 60 minutes during CNO passage. eDREADDs BOLD time courses were also significantly different from Shams and saline CTLs BOLD time courses (from the injection time (15 minutes) until the end of acquisition p< 0.001). iDREADDs time courses were significantly different from Shams’s (p< 0.001 from 25 min on) and the saline CTLs (p< 0.001 at 44 minutes). Finally, Shams and Saline Controls presented no significant difference (p>0.05).

**Figure 3.**
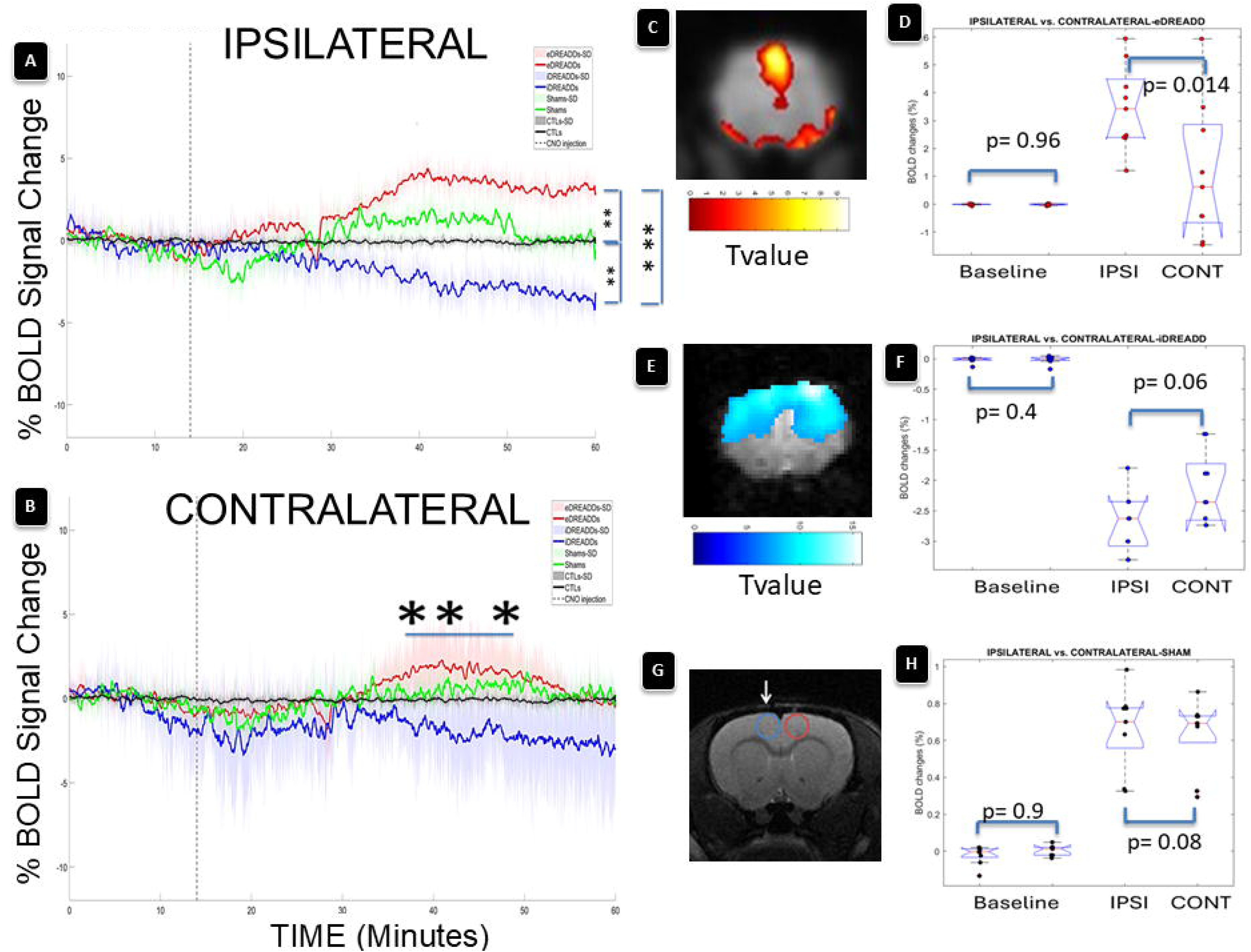
Chemogenetic modulation produces robust and lateralized BOLD responses in the motor cortex. **(A)** Group-averaged ipsilateral BOLD signal time courses during pharmacological fMRI following CNO administration (vertical dashed line). Excitatory DREADD (eDREADD) activation induces a sustained positive BOLD response, whereas inhibitory DREADD (iDREADD) activation elicits a progressive negative BOLD signal compared with Sham and saline control (CTL) animals. Shaded areas represent ± standard deviation (SD). Asterisks indicate time points or intervals with significant group differences as shown in the figure (*p < 0.05, **p < 0.01, ***p < 0.001).**(B)** Contralateral BOLD time courses show attenuated but significant responses in DREADD-expressing animals, with eDREADD animals exhibiting positive and iDREADD animals negative BOLD changes, whereas Sham and CTL groups remain stable. Shaded areas represent ± SD. Horizontal bars with asterisks indicate time windows where group differences reach statistical significance (**p < 0.01, ***p < 0.001), **(C)** statistical parametric maps (t-values) illustrating positive BOLD activation in eDREADD animals. **(D)** Quantification of ipsilateral versus contralateral BOLD signal changes in eDREADD animals at baseline and during activation, showing significant hemispheric lateralization during activation (ipsilateral > contralateral; *p = 0.014), but not at baseline (p = 0.96).**(E).** Corresponding analysis in iDREADD animals shows no significant hemispheric difference at baseline (p = 0.4) and a trend toward lateralization during activation (p = 0.06). **(F).** Negative BOLD activation in iDREADD animals following CNO administration.**(G)** Representative anatomical image showing voxel placement in the right motor cortex. **(H)** Sham animals show no significant hemispheric differences at baseline (p = 0.9) or during activation (p = 0.08). Individual animals are shown with group distributions overlaid. For all groups, *n* = 6-8 rats.

On the contralateral side (Fig.3B), a two-way repeated measures ANOVA revealed a highly significant interaction between Time and Groups, indicating that the effect of Time varied across Groups (F_(33, 288)_ = 38.04, p < 0.0001). This interaction suggests that the temporal pattern of change differed meaningfully among groups. Nevertheless, Time also exhibited a very strong main effect (F_(11, 288)_ = 69.43, p < 0.0001), demonstrating a robust overall influence on the outcome. Groups also showed a significant main effect (F_(3, 288)_ = 661.17, p < 0.0001). Pos-hoc Bonferroni tests indicated significant differences between eDREADDs and iDREADDs contralateral BOLD timecourses in the range 37-54 min (p<0.001). No significant differences were found between BOLD time courses between eDREADDs and Shams or Controls (p>0.05) while iDREADDS BOLD timecourses showed significant differences against Shams and CTLs (p<0.05 from 48 min). No significant difference was found between Shams and CTLs (p>0.05 at all-time points).

### 3.2 Pharmacological fMRI: Excitatory DREADD manipulations produce opposite effects on contra–ipsi BOLD differences over time

Ph-fMRI revealed clear bidirectional modulation of cortical activity: in eDREADD animals, CNO induced strong focal positive BOLD activation in the right ipsilateral MC, extending to homotopic areas (IPSI: +3.5 ± 1.5% vs. CONTRA: +1.1 ± 2.5%; Fig.3C-D). In contrast, interneuron activation DREADD (iDREADD) animals exhibited robust negative BOLD responses (IPSI: −3.3 ± 0.8% vs. CONTRA: −2.7 ± 0.8%; Fig.3E-F), while shams showed no significant changes (IPSI: +0.8 ± 1.5%; CONTRA: +0.7 ± 1.5%; Fig.3H). In Sham rats, the contra–ipsi difference at intercept was small (+0.26) and not statistically significant, indicating roughly equal BOLD responses in contralateral and ipsilateral sides. The difference remained essentially stable over time. eDREADD rats on the other end, showed a small but significant contra > ipsi difference at intercept (+0.30). Over time, the difference decreased slightly (slope = -0.053 per unit time). iDREADD rats exhibited a significant ipsi > contra difference at intercept (-0.92), which was significantly lower than eDREADD at baseline (p < 0.00001). Over time, the difference increased (slope = +0.029), indicating a gradual shift toward contra > ipsi. Between-group comparisons were performed showing that at baseline, iDREADD differed significantly from eDREADD, while Sham did not. The slopes over time were significantly different across all groups, reflecting distinct temporal dynamics: eDREADD decreased slightly, iDREADD increased, and Sham remained flat. Excitatory and interneuron activation DREADD manipulations produce opposite effects on contra–ipsi BOLD differences over time, whereas Sham rats show minimal lateralization.

### 3.3 ^1^H-fMRS revealed no significant neurochemical changes upon CNO activation

Fig. 4A illustrates the VOI position for ^1^H-fMRS acquisitions and Fig. 4B illustrates representative examples of averaged and labeled ^1^H MR spectra acquired in this VOI during baseline and saline infusion in MC in a representative CTL animal pre and post-Saline. Then Shams (Fig. 4C), eDREADDs (Fig. 4D) and iDREADDs (Fig. 4E) time-averaged ^1^H MR spectra are depicted for 2 representative animals per group pre and post-CNO to illustrate the variability across animals and groups. The tCr linewidth ranged between 10.6 ± 0.8 and 12.2 ± 0.2 Hz pre- and post-CNO respectively. The variability across animals and groups was further illustrated in Fig. 4F where the percent change relative to baseline is shown for Glx, GABA and tNAA across groups. Repeated ANOVAs followed by Bonferroni corrections demonstrated no significant differences across groups (p>0.05). Further, the time-averaged metabolite concentrations were calculated during the baseline (0-15 min), during the transition period (16-30 min) and during the active period for CNO (> 30 min) for tNAA, Glx, GABA across groups as well as the SNR at baseline and post-injection (Saline and CNO). No significant concentration changes were found within groups. Only a trend towards decreased tNAA was observed in eDREADDs. Across groups, metabolite concentration and SNR variability was observed attributed to physiological changes during long scanning times (sedation, motion, CNO and saline injections) (Supplementary Figure 1). The mean CRLBs per group as a function of time can be found in the supplementary tables. Metabolite concentration time courses with 5-minute temporal resolution were obtained for Glx (Glu + Gln) (Fig. 5A), GABA (Fig. 5B), tNAA (Fig. 5C), tCr (Fig. 5D) and Asp (Fig. 5E) during CNO uptake in MC for eDREADD and iDREADD, Sham and CTL rats and compared across groups. Time course for the Glx to GABA ratio (Fig. 5F) were also calculated to explore the excitatory to inhibitory balance (E/I). Using a linear mixed model analysis (LMM), we tested for the overall difference in the average metabolite concentrations between the 4 groups across all time points, for the effect of time and for the interaction between group and time. Table 2 summarizes the outcomes of the LMM. Variability across animals was relatively high for some metabolites, particularly tNAA and tCr. Such variability is not unexpected in rodent fMRS studies, where measurements reflect bulk metabolite concentrations within relatively small brain volumes and are influenced by physiological state, anesthesia, and inter-animal differences in viral expression and chemogenetic activation.

**Figure 4.**
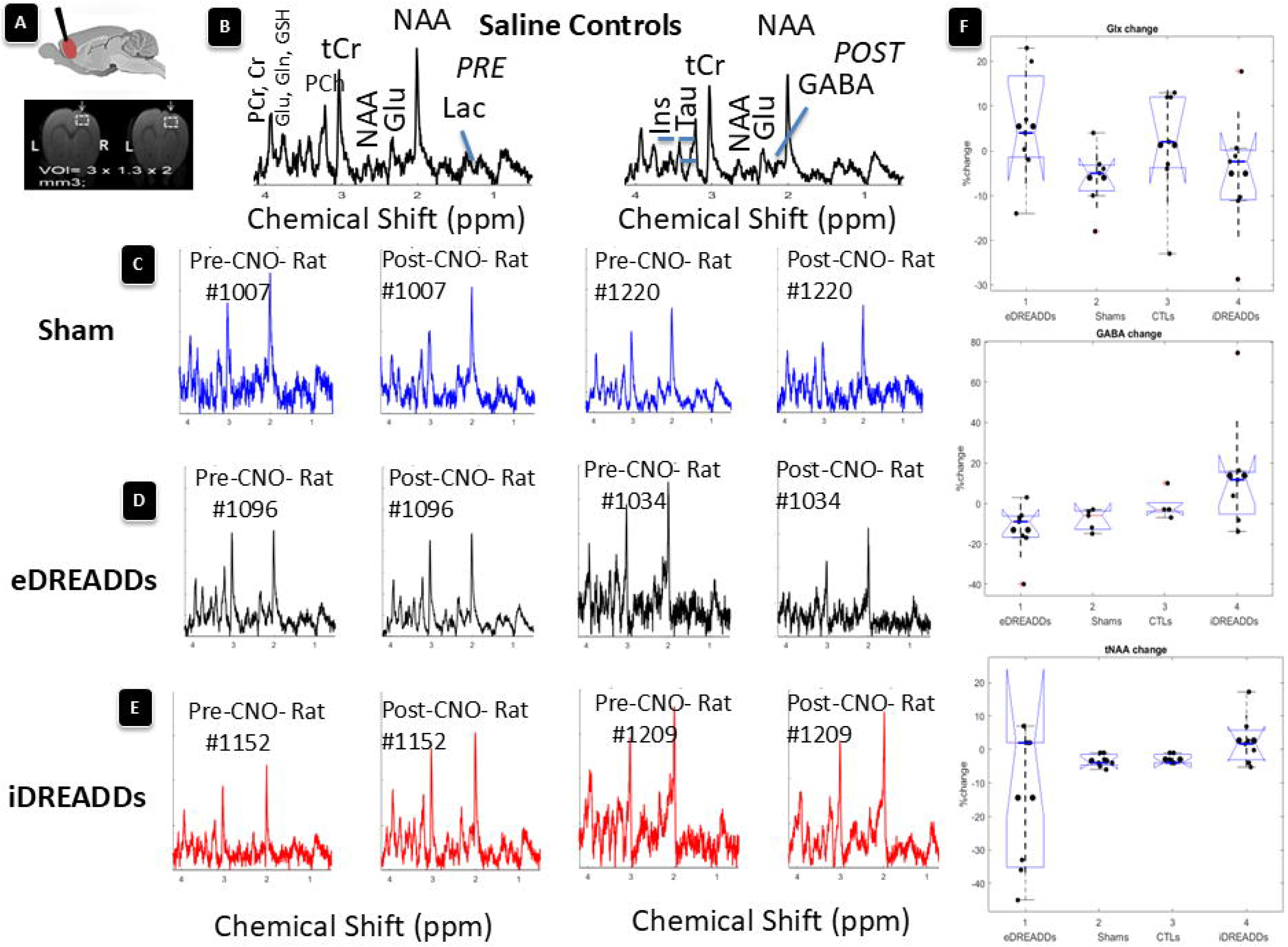
Functional metabolic responses measured with ¹H-fMRS during chemogenetic activation. **(A)** Schematic illustration of the ¹H-fMRS voxel of interest (VOI; 3 × 1.3 × 2 mm³) positioned over the right motor cortex at the site of viral injection, shown on anatomical reference images in coronal orientation. **(B)** Representative time-averaged ¹H MR spectra acquired in saline controls, before (Pre-saline) and during (Post-saline). Major metabolite resonances are labeled, including glutamate/glutamine (Glx), GABA, NAA, creatine compounds, lactate, and choline-containing compounds. Corresponding representative Pre-CNO and Post-CNO spectra acquired in **(C)** Shams **(D)** eDREADD animals and **(E)** iDREADD animals from the same cortical region show comparable spectral profiles before and after CNO administration. Examples for 2 rats per group are shown. Across groups, no obvious qualitative changes in spectral shape or in the relative amplitudes of major metabolite resonances are observed between Pre-CNO and Post-CNO conditions. These representative spectra illustrate the absence of detectable acute metabolic changes at the level of bulk metabolite pools during chemogenetic activation, despite robust BOLD responses observed in parallel fMRI experiments (Fig. 3). All spectra are shown for illustrative purposes; quantitative statistical analyses are reported in subsequent figures. For all groups, *n* = 6- 8 rats.**(F)** The time-averaged % change relative to baseline were calculated for each group and are represented as box-plots with individual points per animal for Glx, GABA and tNAA concentration changes. Repeated ANOVAs followed by Bonferroni corrections showed no significant changes between groups (p>0.05).

**Figure 5.**
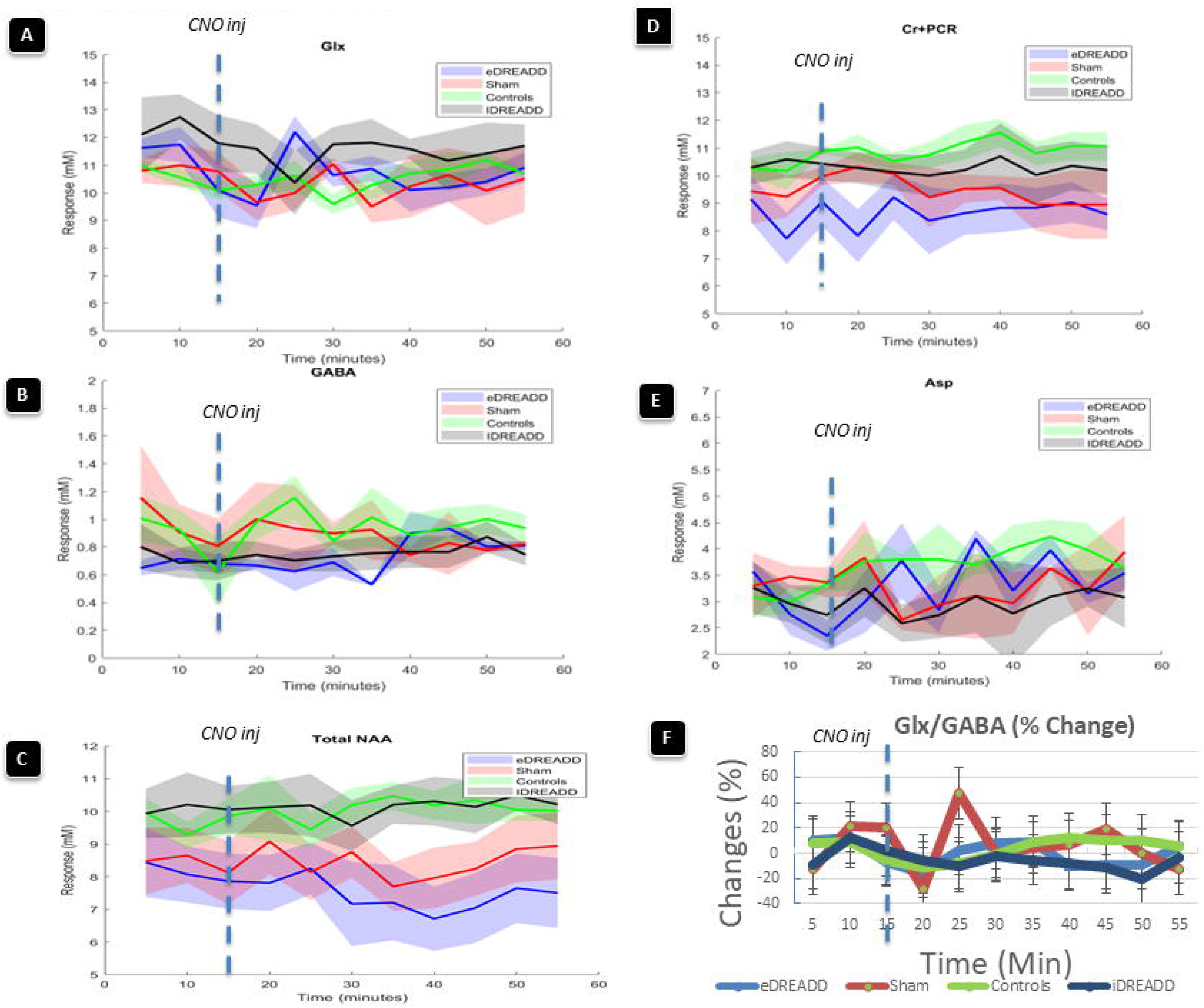
Time-resolved ¹H-fMRS reveals stable bulk metabolite pools during chemogenetic activation (A–E) Group-averaged metabolite concentration time courses acquired from the right motor cortex before and after CNO injection (vertical dashed line). Traces represent group means, and shaded areas indicate ± standard deviation (SD). Panels show: **(A)** glutamate + glutamine (Glx), **(B)** GABA, **(C)** total N-acetylaspartate (tNAA = NAA + NAAG), **(D)** total creatine (Cr + PCr), and **(E)** aspartate (Asp) for excitatory DREADD (eDREADD), inhibitory DREADD (iDREADD), Sham, and saline Control animals.(F) Percent-change time courses of the Glx/GABA ratio (mean ± SD) show modest fluctuations across time without consistent divergence between groups during the expected window of chemogenetic activation.Linear mixed-effects modeling of these time courses (Table 2) revealed significant baseline (intercept) group differences for Glx, GABA, and the Glx/GABA ratio—reflecting chronic metabolic differences between DREADD groups—but no significant acute Group × Time (slope) effects following CNO administration for any metabolite (all slope Cohen’s *d* < 0.1). Minor time-dependent drift effects observed for tNAA and total creatine showed negligible effect sizes and were not specific to DREADD activation.Together, these data demonstrate that, despite robust and opposing BOLD responses (Figure 3), acute chemogenetic modulation does not produce detectable changes in bulk neurometabolite concentrations at 7 T, highlighting a dissociation between hemodynamic and metabolic readouts. For all groups, *n* = 6-8 rats.

**Table 2:** LMM outcomes for raw metabolite concentration time courses: **The Chronic Effect (The Intercepts):** The iDREADDs were in a fundamentally different metabolic state before any CNO was injected. With effect sizes |d|∼ 2.0, this isn’t just a minor statistical fluke; it is a major physiological shift in baseline Glx and GABA, pointing toward a chronic change likely induced by the viral expression itself (e.g., neuroinflammation or altered glial buffering). **The Acute Effect (The Slopes):** The CNO injection, despite successfully triggering a BOLD response did not trigger changes on the bulk metabolite concentrations. The negligible effect sizes associated with acute CNO-induced changes (Cohen’s d < 0.1) indicate that bulk glutamatergic and GABAergic metabolite pools remained remarkably stable despite robust BOLD responses. These findings suggest that acute neurovascular responses can occur in the absence of detectable changes in bulk neurometabolic concentrations, consistent with a degree of neurometabolic dissociation under the present experimental conditions.

| Metabolite | Model Parameter | Comparison/Contrast | P-Value (Holm) | Cohen's d | Biological Interpretation |
| --- | --- | --- | --- | --- | --- |
| **Glx** | Intercept | eDREADD vs. iDREADD | 0.0047 | -2.025 | Baseline difference |
| **Glx** | Slope<br>(15-59 min) | Acute CNO effect | >0.05 | <0.05 | Functional uncoupling |
| **GABA** | Intercept | iDREADD vs. Sham | <0.01 | -1.880 | Baseline difference |
| **GABA** | Slope | Acute CNO effect | >0.05 | <0.03 | Functional uncoupling |
| **Glx/GABA** | Intercept | iDREADD vs. Sham | 0.0088 | 2-342 | Baseline E/I |
| **Glx/GABA** | Slope | Acute CNO effect | >0.05 | <0.06 | Stable E/I ratio |
| **tCr** | Intercept | iDREADDs vs. eDREADDs | 0.142 | -1.256 | Baseline difference |
| **tCr** | Intercept | iDREADDs vs. Sham | 0.626 | 0.652 | Moderate baseline difference |
| **tCr** | Slope | Acute CNO effect | 0.039 | <0.05 | Stability/Minimal drift |
| **tNAA** | Intercept | eDREADDs vs. Saline CTL | p>0.05 | <0.05 | All the groups<br>had similar [tNAA] baseline |
| **tNAA** | Slope | eDREADDs vs. Saline CTL | p=0.048 | <0.05 | Signif. Decreasing [tNAA] over time |

For all the metabolites and the Glx to GABA ratio, the most important finding is the general lack of significant difference in the slopes (Group x time interaction term) after the CNO injection at 15 minutes (post-hoc slope analysis results). No significant differences were observed across groups for Glx, tNAA or tCr. eDREADD activation did not increase the rate of Glx production/consumption compared to saline-CTL and tCr was stable, suggesting CNO did not affect overall cellular energy balance or membrane integrity differently across groups. In addition, the CNO-driven effect did not shift the rate of E/I balance (Excitatory/Inhibitory) differently in DREADD groups although the rate of change of iDREADDs relative to eDREADDs and Shams were close to significance (p=0.09 and p=0.054, respectively). Only GABA concentration time courses showed significant differences between eDREADDs and Shams (p = 0.05) and iDREADDs and Shams (p = 0.009), respectively. The Shams were also initially different from CTL but not after Bonferroni correction.

### 3.4 The Impact on Basal Metabolism

The significant differences at the baseline before CNO were the strongest findings of the present study. These results point toward long-term functional adaptation caused by the DREADD gene expression itself, before the CNO trigger: the basal Glx concentrations in eDREADDs were different from iDREADDs (p =0.004), suggesting chronic differences in glutamatergic tone or basal energy state. The iDREADD group started with significantly lower GABA concentrations than the saline CTL group (p=0.020). This is counter-intuitive and suggests that chronic expression of the iDREADD may lead to a compensatory downregulation of GABA. Hence, the iDREADD group had a significantly higher Glx to GABA ratio (or E/I balance) compared to all other groups. Although iDREADD is designed for inhibition, its baseline E/I balance is shifted toward excitation, likely due to the low basal GABA concentrations. The fact that tCr levels remained stable across all groups throughout the 60 minutes, with no significant slope differences, is also crucial: tCr is primarily an energy buffer. Its stability suggests that while neuronal energy demand is changing (based on BOLD), the overall supply of high-energy phosphates remains adequate and is not being acutely depleted or augmented differently in the DREADD groups.

### 3.5 Effect size analysis of Glx and GABA during baseline and CNO challenge

To complement inferential statistics, effect sizes (Cohen’s D) were computed for Glx and GABA concentration time courses during the baseline period and throughout the CNO challenge. For Glx, very large effect sizes were observed when comparing eDREADD and iDREADD groups, with Cohen’s D exceeding - 2 across the entire time course, including during baseline and persisting throughout CNO administration. The magnitude of this effect remained stable over time and did not show further amplification following CNO injection (Supplementary Fig. 2A). In contrast, comparisons between eDREADD and Sham animals yielded moderate-to-large effect sizes during baseline (Supplementary Fig. 2B), which decreased during the CNO challenge, indicating a relative convergence of Glx levels following CNO administration. Across windows, effect sizes for eDREADD vs Sham were substantially smaller than those observed for eDREADD vs iDREADD. For GABA concentration timecourses, effect sizes were generally smaller than for Glx but followed a similar pattern. eDREADD vs iDREADD comparisons produced consistently larger Cohen’s D values than comparisons involving Sham animals. While modest temporal fluctuations were observed during the CNO challenge, no systematic increase in effect size relative to baseline was detected. Together, these results indicate that the largest group separations in Glx and GABA were present prior to CNO administration, with acute CNO challenge producing limited additional separation at the level of bulk metabolite concentrations, despite robust group differences.

To assess whether the absence of detectable metabolic changes at 7T could be explained by sensitivity limitations rather than biological absence, we performed simulations of Glx and GABA functional time courses under identical chemogenetic perturbations at 7T and 14T. The simulations incorporated realistic pharmacokinetics, compartmental neurometabolic dynamics, and field-strength–dependent noise characteristics, while preserving the same underlying biological effect size across field strengths (See supplementary data for Python codes).

### 3.6 Late CNO/CLZ-driven changes (70 min post-CNO)

A two-way analysis of variance (ANOVA) was performed to assess the effect of acquisition site on neurometabolite concentrations approximately 70 minutes post-CNO administration. The analysis revealed an extremely significant interaction between the Hemisphere and Metabolite factors (F_(55, 408)_ = 17.95, p < 0.0001). This strong interaction demonstrates that the effect of the Hemisphere on metabolite concentration is not uniform; instead, the concentration of any given metabolite in a specific hemisphere is dependent on the type of metabolite being measured. While highly significant main effects were also observed for both Hemisphere (F _(5, 408)_ = 140.06, p < 0.0001) and Metabolite (F_(11, 408)_ = 35.13, p < 0.0001). Consequently, targeted post-hoc Bonferroni comparisons were performed to fully characterize the specific regional differences for each Metabolite-Hemisphere pair, which are presented below:

Within eDREADDs animals, several metabolites exhibited significant hemispheric asymmetries (Table 3). Specifically, GABA and Ins concentrations were significantly reduced in the contralateral hemisphere, whereas GPC+PCh was significantly increased contralaterally, indicating a shift affecting inhibitory tone and membrane turnover. In iDREADDs animals, hemispheric differences were more restricted (Table 3). A significant contralateral increase in GPC+PCh was observed, suggesting altered high-energy phosphate metabolism. In contrast, Sham animals displayed minimal hemispheric differences (Table 3), with only a limited number of metabolites reaching significance. Notably, Asp (+49.8%, p < 0.001) was increased contralaterally, confirming an overall symmetric metabolic profile in the absence of DREADDs expression. Between-group comparisons revealed marked differences in the extent and hemispheric distribution of metabolic alterations between eDREADDs and iDREADDs animals. Direct comparison between eDREADDs and iDREADDs animals revealed robust and widespread metabolic differences on both hemispheres (Table 4). These differences were markedly more extensive on the contralateral side. Across hemispheres, iDREADDs animals showed higher levels of high-energy phosphates and amino-acid–related metabolites relative to eDREADDs (Table 4). In eDREADDs rats, metabolic differences relative to Sham animals were limited on the ipsilateral side, with significant effects restricted to a small subset of metabolites (notably GABA, Ins, and GPC+PCh) (See Supplementary Table 1). In contrast, comparisons on the contralateral side revealed a larger number of significant alterations involving amino acids, energy-related metabolites, and phosphocholine compounds. In iDREADDs animals, comparisons with Sham animals revealed significant alterations across a broader range of metabolites on both hemispheres. These differences predominantly involved energy-related compounds such as glucose, and amino-acid–related metabolites (Asp, Glu+Gln, Tau, NAA+NAAG), indicating a more widespread metabolic divergence from Sham controls (See Supplementary Table 1).

**Table 3:** Hemispheric Differences Across Groups (70 min post-CNO): Comparison between contralateral and ipsilateral metabolite concentrations. p-values are Bonferroni-corrected (n = 12 comparisons per group).

| Metabolite | eDREADDs $\Delta$ (%) | SD | p | Sig | d | iDREADDs $\Delta$ (%) | SD | p | Sig | d | Sham $\Delta$ (%) | SD | p | Sig | d |
| --- | --- | --- | --- | --- | --- | --- | --- | --- | --- | --- | --- | --- | --- | --- | --- |
| **Asp** | -15.8 | 24.4 | >0.05 | ns | 0.65 | -1.78 | 13.6 | 0.05 | ns | -0.18 | +49.8 | 8.9 | 0.001 | *** | 7.9 |
| **GABA** | -18.0 | 11.1 | <0.05 | * | 0.85 | +10.0 | 9.9 | 0.05 | ns | 1.43 | +11.5 | 1.6 | 0.05 | ns | 10.0 |
| **Glc** | +7.4 | 19.2 | >0.05 | ns | 0.38 | +6.32 | 7.28 | 0.05 | ns | 1.23 | +6.4 | 3.0 | 0.05 | ns | 3.0 |
| **Ins** | -29.5 | 11.4 | <0.001 | *** | 2.58 | +4.69 | 7.81 | 0.05 | ns | 0.85 | +8.3 | 2.8 | 0.05 | ns | 4.2 |
| **Tau** | -17.0 | 10.6 | <0.05 | * | 0.83 | +4.58 | 9.9 | 0.05 | ns | 0.65 | -11.3 | 3.7 | 0.05 | ns | -4.4 |
| **GPC+PCh** | +20.2 | 13.3 | <0.01 | ** | 0.87 | +19.8 | 5.62 | 0.01 | ** | 1.79 | -16.0 | 2.3 | 0.05 | ns | -10.0 |
| **NAA+NAAG** | -7.6 | 8.1 | >0.05 | ns | 0.94 | +0.72 | 7.07 | 0.05 | ns | 0.14 | +14.4 | 2.1 | 0.05 | ns | 9.8 |
| **Glu+Gln** | -4.1 | 17.0 | >0.05 | ns | 0.24 | +10.84 | 4.24 | 0.05 | ns | 3.61 | +3.0 | 1.0 | 0.05 | ns | 4.2 |
| **PCr +Cr** | -4.8 | 6.0 | >0.05 | ns | 0.84 | -4.8 | 4.0 | 0.05 | ns | -1.65 | +9.8 | 1.7 | 0.05 | ns | 8.2 |

**Table 4:**
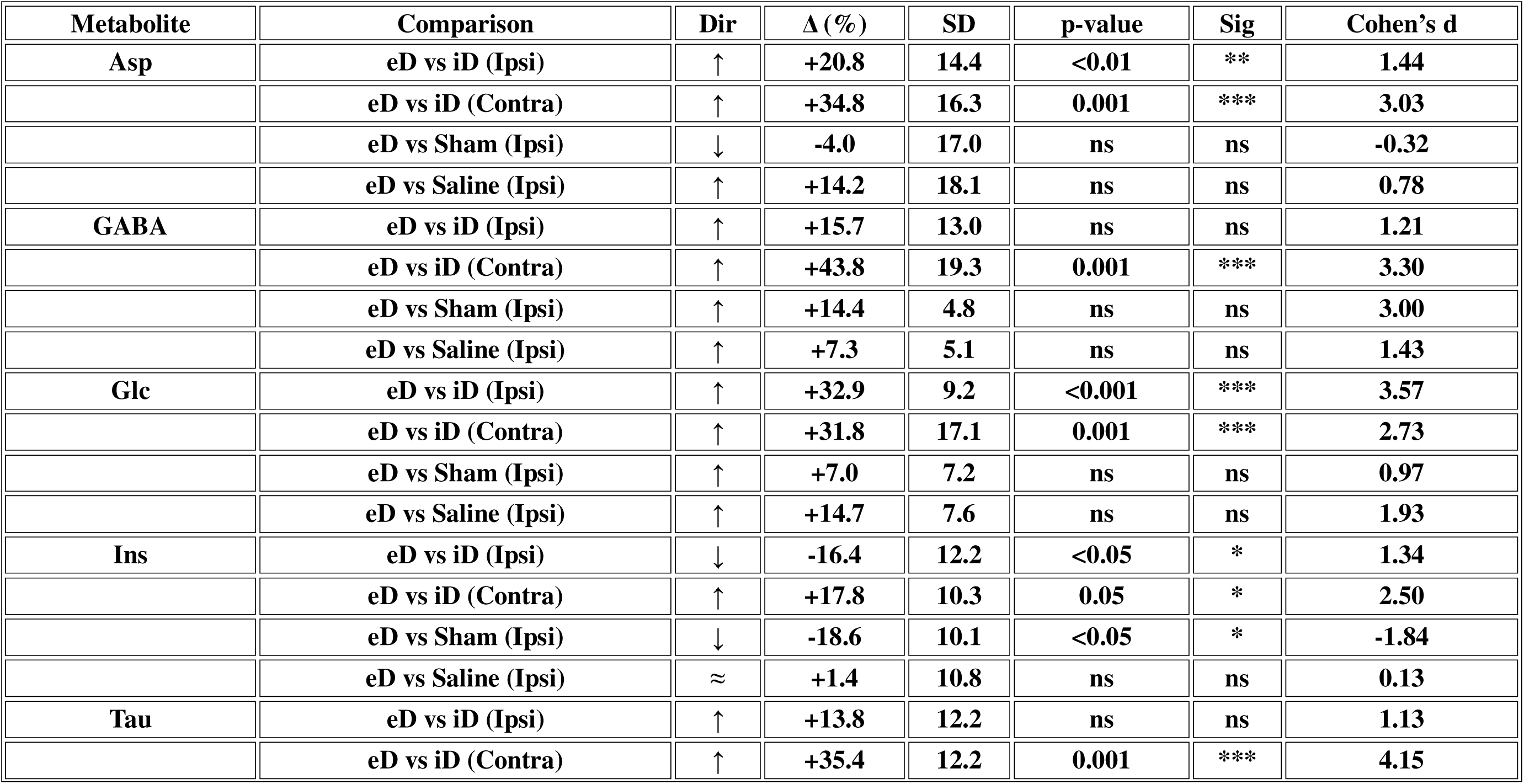
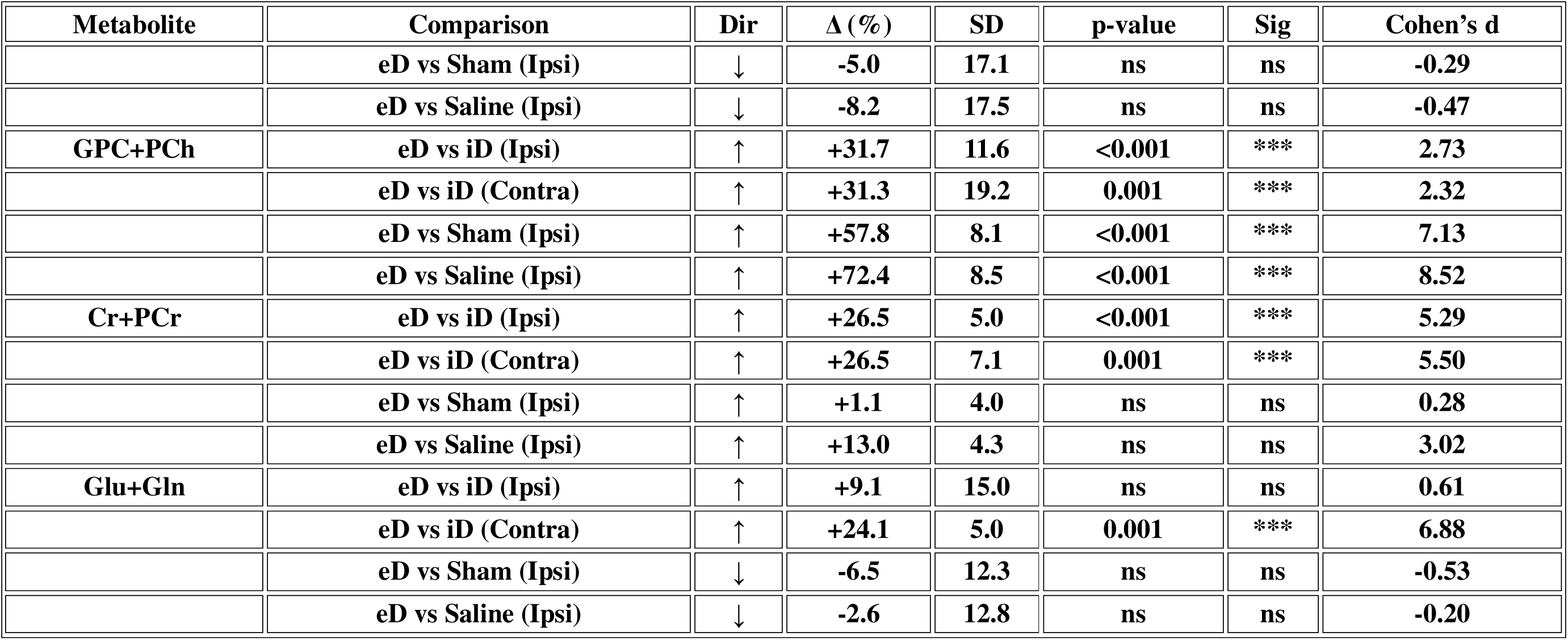
Consolidated Late-Point Metabolic Comparisons (n = 44 comparisons) Note: p-values are Bonferroni-corrected for 44 comparisons. Direction (Dir) and percentage change (Δ %) refer to the first group relative to the second. eDREADDs (eD), iDREADDS (iD)

In Saline control animals, hemispheric comparison performed approximately 70 min after injection revealed significant differences only for Glc, Asp (p<0.05) and total choline (GPC+PCh, p<0.05)) after Bonferroni correction, with no other metabolites affected. A similar restricted pattern was observed when Saline controls were compared with Sham animals on either hemisphere (See Supplementary Table 1). Thus, under control conditions, delayed static MRS reveals only minimal hemispheric metabolic asymmetry, indicating that the widespread alterations observed in DREADD-expressing animals are not attributable to injection-related or time-dependent effects. Relative to Saline controls, DREADD-expressing animals exhibited marked and asymmetric metabolic alterations at the delayed static MRS time point (∼70 min post-injection). In eDREADDs animals, the ipsilateral hemisphere differed from Saline animals (both ipsilateral and contralateral) primarily for total choline (GPC+PCh), which was significantly altered on both sides. In contrast, the contralateral hemisphere of eDREADDs animals showed broader deviations: compared with Saline-Ipsi, significant differences were observed for Asp, GABA, Ins, and total choline, whereas comparison with Saline-Contra revealed a significant effect only for GABA (See Supplementary Table 1). In iDREADDs animals, the ipsilateral hemisphere differed from Saline-Ipsi for Asp, Ins, and Tau, and from Saline-Contra for a wider panel of metabolites, including Asp, Glc, Tau, total NAA, and tCr. The contralateral hemisphere of iDREADDs animals exhibited the most extensive divergence, differing significantly from both Saline-Ipsi and Saline-Contra across the majority of measured metabolites. Thus, while Saline animals displayed only minimal hemispheric metabolic asymmetries, DREADD-expressing animals showed robust and hemisphere-dependent deviations, with iDREADDs animals exhibiting the broadest and most pronounced late metabolic alterations (See Supplementary Table 1). The mean CRLBs per group can be found in the supplementary tables ‘document.

## 4. Discussion

The exact metabolic cost of DREADD-mediated circuit manipulation, particularly with respect to acute energy demand, neurotransmitter turnover, and neurovascular signaling, remains poorly understood. To address this gap, we combined longitudinal BOLD fMRI and ^1^H-fMRS in the motor cortex of sedated rats using a neuro-preserving imaging protocol [16]. Rather than attempting to isolate individual cellular contributions, we exploited the contrasting network-level consequences of chemogenetic recruitment of broad neuronal populations and preferential recruitment of inhibitory interneuron networks. This approach enabled us to investigate how distinct patterns of circuit engagement influence the relationship between neuronal activity, metabolism, and vascular responses.

A major finding of the present study is the robust dissociation between hemodynamic and neurochemical readouts following chemogenetic modulation of cortical circuits. Whereas these findings could straightaway be ascribed to sensitivity issues of ^1^H-fMRS at a preclinical field strength of 7T [31,32], we also measured and detected significant metabolic interhemispheric changes at a later timepoint (70-min post-CNO) within the same animals. Although the detected changes may rather be the consequences of the CNO to clozapine conversion [33, 34](discussed below), they also show that sensitivity limitations may not explain our findings. Moreover, the comparison between 7 T and 14 T suggests that improved spectral resolution and signal-to-noise ratio at higher field strengths may enhance the detectability of subtle neurometabolic changes. However, the absence of significant acute fMRS responses at 7 T cannot be attributed solely to limited sensitivity. Alternative explanations include biological buffering of neurotransmitter pools, compartmental averaging effects, or temporal dissociation between neuronal activity and measurable changes in bulk metabolite concentrations. Further supporting this, an earlier fMRSI study [35] also demonstrated a coupling mismatch between energy consumption and supply.

In the present study, chemogenetic recruitment of broad neuronal populations elicited pronounced positive BOLD responses, whereas preferential recruitment of inhibitory interneuron networks produced marked negative BOLD responses. Despite these opposing hemodynamic effects, neither manipulation induced proportional or consistent alterations in glutamatergic or GABAergic metabolite pools measured by ^1^H-fMRS during the CNO challenge. These findings demonstrate that substantial vascular responses can emerge in the absence of measurable changes in bulk neurotransmitter concentrations and support direct experimental evidence for the existence of neurometabolic dissociation under conditions of externally driven circuit modulation. The contrasting responses observed between broad neuronal recruitment and interneuron-enriched recruitment further suggest that the magnitude and direction of the BOLD response depend strongly on the cellular composition of the engaged network [6,7]. Conceptually, broad neuronal recruitment reflects the combined influence of excitatory and inhibitory neuronal populations, whereas interneuron-enriched recruitment preferentially emphasizes inhibitory network contributions [15, 36]. Under this framework, the difference between these conditions may approximate the net excitatory component of the neurovascular response. However, because cortical circuits are highly recurrent and nonlinear, this interpretation should be considered a systems-level conceptual model rather than a direct measure of excitatory neuronal activity. Importantly, the absence of detectable neurotransmitter changes does not necessarily imply the absence of altered neurotransmission. Functional MRS measures changes in relatively large metabolite reservoirs [8,37,38], whereas neuronal communication relies on rapid neurotransmitter release, uptake, recycling, and compartmental shifts [39]. Consequently, substantial changes in neurotransmitter flux and synaptic activity may occur without producing measurable alterations in steady-state glutamate or GABA concentrations. The present findings therefore suggest that the observed BOLD responses are not simply determined by bulk neurotransmitter pool size but instead reflect broader neurovascular processes. One potential explanation for this dissociation is the differential engagement of the neurovascular unit. Astrocytes, pericytes, endothelial cells, and vascular smooth muscle cells collectively contribute to the translation of neuronal activity into local hemodynamic responses[40, 41,42].Astrocytes are of particular interest because they respond to both glutamatergic and GABAergic signaling and can regulate cerebral blood flow through calcium-dependent release of vasoactive mediators [43]. Under this framework, distinct patterns of neuronal recruitment may generate markedly different vascular responses despite producing minimal changes in measurable neurotransmitter concentrations. The positive BOLD responses associated with broad neuronal recruitment and the negative responses observed following preferential inhibitory network recruitment may therefore reflect differential engagement of neurovascular signaling pathways rather than differences in steady-state neurotransmitter levels alone. This interpretation aligns with a growing body of literature challenging the assumption of a tight and linear relationship between neuronal activity, metabolism, and hemodynamics. Pereira et al. [44] recently demonstrated a dissociation between metabolic and hemodynamic responses during chemogenetic manipulation of astrocytes in the mouse visual cortex. Previous rodent studies have similarly shown that BOLD, cerebral blood flow, and cerebral blood volume responses do not always scale proportionally with neuronal or metabolic activity. For example, hippocampal seizures can generate large increases in neuronal firing and perfusion while producing paradoxical negative BOLD responses [45], suggesting that oxygen consumption may outpace vascular delivery. Likewise, perforant-path stimulation induces prolonged hemodynamic depressions despite persistent after-discharges [46], indicating the existence of multiple and potentially independent neurovascular coupling mechanisms. Pharmacological studies have further demonstrated that vascular responses depend strongly on the receptor systems engaged, with NMDA- and AMPA-mediated signaling producing distinct hemodynamic consequences despite ongoing neuronal excitation [47]. Together, these findings indicate that coupling between vascular signals and glutamate or GABA metabolism is conditional, context-dependent, and influenced by multiple cellular pathways. In humans, uncoupling is implicated in aging and neurodegenerative disorders such as Alzheimer’s disease, where pericyte loss, astrocytic dysfunction, and oxidative stress impair vascular dilation and metabolic efficiency [48]. This has implications for the interpretation of combined fMRI–fMRS studies, particularly those aiming to infer neurotransmitter dynamics from BOLD signals [9, 35, 49–51]. The present results argue that caution is warranted when equating hemodynamic changes with neurotransmitter metabolism, especially in experimental paradigms involving neuromodulatory or chemogenetic interventions. The convergence of evidence across different cell types (neurons and astrocytes), cortical regions (motor and visual cortex), species, and experimental paradigms suggest that neurometabolic dissociation may represent a fundamental biological feature of externally driven circuit perturbations rather than a methodological artifact. Because DREADD-mediated modulation produces sustained cellular recruitment that differs from endogenous activity patterns, vascular responses may be strongly influenced by downstream vasoactive signaling pathways without necessarily requiring large-scale shifts in total neurotransmitter reservoirs. Collectively, our results support the view that hemodynamic signals emerge from interactions among multiple components of the neurovascular unit and that BOLD and fMRS measurements capture complementary but distinct aspects of circuit function.

An important observation emerging from the present dataset is that substantial neurometabolic differences were already present at baseline between eDREADDs- and iDREADDs-expressing animals, particularly for Glx and related metabolites, with very large effect sizes (Cohen’s *d* > 2). These differences precede ligand administration and therefore cannot be attributed to acute chemogenetic activation, CNO pharmacodynamics, or clozapine back-metabolism [33, 34]. Instead, they suggest that chronic expression of the constructs may contribute to distinct neurometabolic states.. Such baseline divergence is biologically plausible. Sustained expression of excitatory versus inhibitory DREADDs can alter intrinsic excitability, synaptic balance, and network tone over weeks, even in the absence of ligand [33, 34, 52–53]. Chronic perturbations of neuronal excitability are known to trigger homeostatic plasticity, synaptic scaling, and compensatory changes in excitation–inhibition balance [53–55], processes that are metabolically costly and can reshape bulk pools of amino acids and high-energy phosphates. In parallel, long-term viral transduction itself can modify cellular physiology, glial engagement, and local metabolic demand [56]. In this context, ¹H-MRS captures a trait-like metabolic phenotype associated with circuit identity rather than a state-like response to acute activation [26]. This interpretation reframes the largely negative dynamic ^1^H-fMRS findings: the dominant chemogenetic effect detectable at 7 T is not a transient modulation during the CNO challenge, but a stable reconfiguration of neurometabolic baseline. Acute BOLD responses then unfold on top of these pre-existing metabolic landscapes. The dissociation between large baseline differences and minimal dynamic changes suggests that chemogenetic constructs primarily bias long-term metabolic organization, while short-term vascular and electrophysiological responses are mediated through mechanisms that do not require proportional shifts in bulk neurotransmitter pools [57–59]. Importantly, the use of a broad hSyn promoter for excitatory DREADDs may have further contributed to this effect by engaging heterogeneous neuronal populations and inducing diffuse homeostatic adaptations [15, 60]. Future work using cell-type–specific promoters and longitudinal pre- and post-transduction MRS will be necessary to disentangle viral-expression effects from ligand-driven neuromodulation.

In contrast to the acute ^1^H-fMRS results, interhemispheric metabolic differences were observed approximately 70 minutes after CNO administration, when measurements were performed using static ^1^H-MRS rather than ^1^H-fMRS. These late effects were not temporally aligned with the BOLD responses and therefore likely reflect secondary or delayed metabolic processes rather than direct consequences of acute circuit activation. Crucially, the delayed emergence of hemispheric asymmetries suggests that these metabolic changes are not causally linked to the BOLD dynamics observed earlier, reinforcing the interpretation of neurometabolic dissociation at acute timescales. A critical consideration in interpreting the delayed metabolic effects is the known back-metabolism of CNO to clozapine in rodents [33,34]. Clozapine exhibits high affinity for multiple endogenous receptors and can exert widespread neuromodulatory and metabolic effects independent of DREADDs activation [61–63]. The timing of the observed interhemispheric differences is compatible with a scenario in which clozapine accumulation contributes to delayed metabolic alterations. While the present study was not designed to directly dissociate CNO- from clozapine-mediated effects, the temporal dissociation between BOLD, ^1^H-fMRS, and static MRS findings strongly suggests that late metabolic changes should be interpreted with caution. For this reason, these results are best viewed as hypothesis-generating rather than definitive evidence of “chemogenetically” driven metabolic reorganization.The observation that delayed metabolic changes were often more pronounced contralaterally further supports the idea that these effects arise from distributed network processes or systemic influences, rather than localized neuronal activation. Importantly, Sham animals did not exhibit comparable hemispheric asymmetries, indicating that these effects are not attributable to measurement bias or baseline lateralization. However, given their delayed onset and the pharmacokinetic considerations discussed above, these interhemispheric differences likely reflect indirect consequences of chemogenetic manipulation and/or clozapine exposure, rather than primary neurometabolic correlates of BOLD activation.

As multimodal neuroscience continues to develop, addressing the underlying biophysical relationships between imaging modalities may prove just as important as advancing the measurement technologies themselves. To move beyond purely observational neuroimaging, the field must leverage causal perturbation methods, such as chemogenetics, optogenetics, transcranial magnetic stimulation (TMS), transcranial focused ultrasound (TFUS), and genetically encoded biosensors [64] specifically to address how these independent axes of neurovascular and neurometabolic coupling operate.

## 5. Conclusion

In this study, chemogenetic excitation elicited large, opposing BOLD responses, yet these rapid hemodynamic changes were not accompanied by contemporaneous alterations in glutamate or GABA concentrations measured with ^1^H-fMRS at 7T. Rather than undermining the utility of fMRS, these results highlight the complementary information provided by hemodynamic and neurochemical imaging modalities. The absence of detectable acute metabolite changes despite robust BOLD responses suggests that neurovascular and neurometabolic readouts may not always evolve in parallel, cautioning against directly inferring acute neurotransmitter turnover from BOLD dynamics alone. Our work highlights that neurovascular and neurometabolic readouts can be experimentally dissociated. It provides a critical roadmap that motivates future multimodal studies to move beyond static pool sizes at standard field strengths, leveraging next-generation stable ligands, ultra-high fields (> 14T), and dynamic flux kinetics to truly map out the independent pathways governing neurovascular and neurometabolic coupling.

## Supporting information

Simulations

## Abbreviations

Ala: Alanine
Asc: Ascorbate
Asp: Aspartate
ATP: Adenosine Triphosphate
BOLD: Blood Oxygen Level dependent
CC: Corpus Callosum
CNO: Clozapine-N-oxide
CRLB: Cramér Rao Lower Bounds
Cr: Creatine
DREADDs: Designer Receptors Activated Only by Designer Drugs
eDREADDS: excitatory DREADDs;
iDREADDs: Interneuron DREADDs
fid: free induction decay
fMRI: functional magnetic resonance imaging
fMRS: functional magnetic resonance spectroscopy
GABA: γ-aminobutyric acid
Glc: Glucose
Glu: Glutamate
Gln: Glutamine
Glx: Glutamate + Glutamine
GPC: Glycerophosphocholine
GRE-EPI: Gradient Echo Echo Planar Imaging
HR: Heart Rate
Ins: Myo-Inositol
LCModel: Linear Combination of Model
Mac: Macromolecules
MED-ISO: Medetomidine-isoflurane sedation
MC: Motor Cortex
MS: Multiple Sclerosis
NAA: N.Acetyl-Aspartate
NAAG: N-acetyl-Aspartyl Glutamic acid
PBS: Phosphate Buffered Saline
PCh: Phosphocholine
PCr: Phosphocreatine
PFA: Paraformaldehyde
Ph-fMRI: Pharmacological MRI
Tau: Taurine
tCr: Total Creatine
tNAA: total N-Acetyl-Aspartate
Scyllo: Scyllo-Inositol
SNR: Signal to Noise Ratio
SpO2: Partial Pressure of Oxygen

## Author Contributions

The project was conceived and designed by NJ. The experiments were performed by NJ. The manuscript was drafted and edited by FAV and NJ. Writing–review and editing were done by FAV and NJ. Resources and supervision were provided by NJ. All authors have read and agreed to this version of the manuscript.

## Acknowledgments

This work was supported by the Lundbeck Foundation (Experiment grant, grant nr. R370-2021-402) to NJ. Animal viral injections, histology and immunohistochemistry were performed by technicians at the DRCMR. We thank the support of the Leenards -Jeantet laboratory for functional et metabolic imaging (LIFMET) and help from colleagues to sort data and quantify immunohistochemical data.

## Ethics Statement

All experiments were approved by the Danish Animal Experiments Inspectorate (2018-15-0201-01551) and follow the European Communities Council Directive (2010/63/EU).

## Conflicts of Interest

Authors declare no conflicts of interest.

## Data Availability Statement

Data will be made available upon reasonable request.

## Notes

### Competing Interest Statement

The authors have declared no competing interest.

## References

1. Andrea Giorgi, Sara Migliarini, Alberto Galbusera, Giacomo Maddaloni, Maddalena Mereu, Giulia Margiani, Marta Gritti, Silvia Landi, Francesco Trovato, Sine Mandrup Bertozzi, Andrea Armirotti, Gian Michele Ratto, Maria Antonietta De Luca, Raffaella Tonini, Alessandro Gozzi, Massimo Pasqualetti. Brain-wide Mapping of Endogenous Serotonergic Transmission via Chemogenetic fMRI. Cell Rep. 2017 Oct 24;21(4):910–918. doi: 10.1016/j.celrep.2017.09.087.

2. Akihiko Ozawa, Hiroyuki Arakawa. Chemogenetics drives paradigm change in the investigation of behavioral circuits and neural mechanisms underlying drug action. Behav Brain Res. 2021 May 21:406:113234. doi: 10.1016/j.bbr.2021.113234.

3. Rocchi F, Canella C, Noei S, Gutierrez-Barragan D, Coletta L, Galbusera A, Stuefer A, Vassanelli S, Pasqualetti M, Iurilli G, Panzeri S, Gozzi A. Increased fMRI connectivity upon chemogenetic inhibition of the mouse prefrontal cortex.Nat Commun. 2022 Feb 25;13(1):1056. doi: 10.1038/s41467-022-28591-3

4. Markicevic M, Fulcher BD, Lewis C, Helmchen F, Rudin M, Zerbi V, Wenderoth N. Cortical Excitation:Inhibition Imbalance Causes Abnormal Brain Network Dynamics as Observed in Neurodevelopmental Disorders.Cereb Cortex. 2020 Jul 30;30(9):4922–4937. doi: 10.1093/cercor/bhaa084.

5. Cornelia Helbing, Marta Brocka, Alberto Arboit, Michael T Lippert, Frank Angenstein. Chemogenetic inhibition of dopaminergic neurons reduces stimulus-induced dopamine release, thereby altering the hemodynamic response function in the prefrontal cortex. Imaging Neurosci (Camb). 2024 Jun 21;2:imag-2-00200. doi: 10.1162/imag_a_00200

6. Gauthier CJ, Fan AP. BOLD signal physiology: Models and applications. Neuroimage. 2019 Feb 15;187:116–127. doi: 10.1016/j.neuroimage.2018.03.018. Epub 2018 Mar 13. PMID: 29544818.

7. Uludağ K. Physiological modeling of the BOLD signal and implications for effective connectivity: A primer. Neuroimage. 2023 Aug 15;277:120249. doi: 10.1016/j.neuroimage.2023.120249. Epub 2023 Jun 24. PMID: 37356779.

8. Just N. Proton functional magnetic resonance spectroscopy in rodents. NMR Biomed. 2021 May;34(5): e4254. doi: 10.1002/nbm.4254.

9. Just N, Faber C.J Probing activation-induced neurochemical changes using optogenetics combined with functional magnetic resonance spectroscopy: a feasibility study in the rat primary somatosensory cortex. Neurochem. 2019 Aug;150(4):402–419. doi: 10.1111/jnc.14799.

10. Bednařík P, Tkáč I, Giove F, DiNuzzo M, Deelchand DK, Emir UE, Eberly LE, Mangia S. Neurochemical and BOLD responses during neuronal activation measured in the human visual cortex at 7 Tesla. J Cereb Blood Flow Metab. 2015 Mar 31;35(4):601–10. doi: 10.1038/jcbfm.2014.233.

11. Duanghathai Pasanta, Jason L He, Talitha Ford, Georg Oeltzschner, David J Lythgoe, Nicolaas A Puts. Functional MRS studies of GABA and glutamate/Glx - A systematic review and meta-analysis. Neurosci Biobehav Rev. 2023 Jan:144:104940. doi: 10.1016/j.neubiorev.2022.104940. Epub 2022 Nov 2.

12. Alexander R Craven, Gerard Dwyer, Lars Ersland, Katarzyna Kazimierczak, Ralph Noeske, Lydia Brunvoll Sandøy, Erik Johnsen, Kenneth Hugdahl. GABA, glutamatergic dynamics and BOLD contrast assessed concurrently using functional MRS during a cognitive task.NMR Biomed. 2024 Mar;37(3):e5065. doi: 10.1002/nbm.5065. Epub 2023 Oct 28.

13. Takado Y, Takuwa H, Sampei K, Urushihata T, Takahashi M, Shimojo M, Uchida S, Nitta N, Shibata S, Nagashima K, Ochi Y, Ono M, Maeda J, Tomita Y, Sahara N, Near J, Aoki I, Shibata K, Higuchi M. MRS-measured glutamate versus GABA reflects excitatory versus inhibitory neural activities in awake mice. J Cereb Blood Flow Metab. 2022 Jan;42(1):197–212. doi: 10.1177/0271678X211045449. Epub 2021 Sep 13. PMID: 34515548; PMCID: PMC8721779.

14. Pasanta D, He JL, Ford T, Oeltzschner G, Lythgoe DJ, Puts NA. Functional MRS studies of GABA and glutamate/Glx - A systematic review and meta-analysis. Neurosci Biobehav Rev. 2023 Jan;144:104940. doi: 10.1016/j.neubiorev.2022.104940. Epub 2022 Nov 2. PMID: 36332780; PMCID: PMC9846867.

15. Dimidschstein J, Chen Q, Tremblay R, Rogers SL, Saldi GA, Guo L, Xu Q, Liu R, Lu C, Chu J, Grimley JS, Krostag AR, Kaykas A, Avery MC, Rashid MS, Baek M, Jacob AL, Smith GB, Wilson DE, Kosche G, Kruglikov I, Rusielewicz T, Kotak VC, Mowery TM, Anderson SA, Callaway EM, Dasen JS, Fitzpatrick D, Fossati V, Long MA, Noggle S, Reynolds JH, Sanes DH, Rudy B, Feng G, Fishell G. A viral strategy for targeting and manipulating interneurons across vertebrate species. Nat Neurosci. 2016 Dec;19(12):1743–1749. doi: 10.1038/nn.4430. Epub 2016 Oct 31. Update in: Nat Neurosci. 2017 Jun 27;20(7):1033. doi: 10.1038/nn0717-1033d. Erratum in: Nat Neurosci. 2017 Jun 27;20(7):1033. doi: 10.1038/nn0717-1033c. PMID: 27798629; PMCID: PMC5348112.

16. Just N, Hoehn M. To intubate or not? Balancing anesthesia in rodent fMRI: strategies to mitigate confounding effects. Cereb Cortex. 2025 Jan 6:bhae499. doi: 10.1093/cercor/bhae499.

17. Barrière DA, Magalhães R, Novais A, Marques P, Selingue E, Geffroy F, Marques F, Cerqueira J, Sousa JC, Boumezbeur F, Bottlaender M, Jay TM, Cachia A, Sousa N, Mériaux S. The SIGMA rat brain templates and atlases for multimodal MRI data analysis and visualization. Nat Commun. 2019 Dec 13;10(1):5699. doi: 10.1038/s41467-019-13575-7. PMID: 31836716; PMCID: PMC6911097.

18. Klomp A, Tremoleda JL, Wylezinska M, Nederveen AJ, Feenstra M, Gsell W, Reneman L. Lasting effects of chronic fluoxetine treatment on the late developing rat brain: age-dependent changes in the serotonergic neurotransmitter system assessed by pharmacological MRI.Neuroimage. 2012 Jan 2;59(1):218–26. doi: 10.1016/j.neuroimage.2011.07.082.

19. Schrantee A, Tremoleda JL, Wylezinska-Arridge M, Bouet V, Hesseling P, Meerhoff GF, de Bruin KM, Koeleman J, Freret T, Boulouard M, Desfosses E, Galineau L, Gozzi A, Dauphin F, Gsell W, Booij J, Lucassen PJ, Reneman L. Repeated dexamphetamine treatment alters the dopaminergic system and increases the phMRI response to methylphenidate.PLoS One. 2017 Feb 27;12(2):e0172776. doi: 10.1371/journal.pone.0172776.

20. Tkac I, Starcuk Z, Choi IY, Gruetter R. In vivo 1H NMR spectroscopy of rat brain at 1 ms echo time. Magn Reson Med 1999; 41:649 – 656. 17.

21. Simpson R, Devenyi GA, Jezzard P, Hennessy TJ, Near J. Advanced processing and simulation of MRS data using the FID appliance (FID-A)-An open source, MATLAB-based toolkit. Magn Reson Med. 2017 Jan;77(1):23–33. doi: 10.1002/mrm.26091.

22. Provencher SW. Automatic quantitation of localized in vivo 1H spectra with LCModel. NMR Biomed. 2001 Jun;14(4):260–4. doi: 10.1002/nbm.698.

23. Hyder F, Patel AB, Gjedde A, Rothman DL, Behar KL, Shulman RG. Neuronal-glial glucose oxidation and glutamatergic-GABAergic function. J Cereb Blood Flow Metab. 2006 Jul;26(7):865–77. doi: 10.1038/sj.jcbfm.9600263. Epub 2006 Jan 11.PMID: 16407855.

24. Gruetter R. In vivo 13C NMR studies of compartmentalized cerebral carbohydrate metabolism. Neurochem Int. 2002 Aug-Sep;41(2-3):143–54. doi: 10.1016/s0197-0186(02)00034-7. PMID: 12020614.

25. Tkác I, Oz G, Adriany G, Uğurbil K, Gruetter R. In vivo 1H NMR spectroscopy of the human brain at high magnetic fields: metabolite quantification at 4T vs. 7T. Magn Reson Med. 2009 Oct;62(4):868–79. doi: 10.1002/mrm.22086. PMID: 19591201; PMCID: PMC2843548.

26. Rae CD. A guide to the metabolic pathways and function of metabolites observed in human brain 1H magnetic resonance spectra. Neurochem Res. 2014 Jan;39(1):1–36. doi: 10.1007/s11064-013-1199-5. Epub 2013 Nov 21. PMID: 24258018.26. Near J. et al., *NMR in Biomedicine*, 2013 – fMRS methodology and limitations

27. Gabrielsson J, Meibohm B, Weiner D. Pattern Recognition in Pharmacokinetic Data Analysis. AAPS J. 2016 Jan;18(1):47–63. doi: 10.1208/s12248-015-9817-6. Epub 2015 Sep 3. PMID: 26338231; PMCID: PMC4706292.

28. Near J, Edden R, Evans CJ, Paquin R, Harris A, Jezzard P. Frequency and phase drift correction of magnetic resonance spectroscopy data by spectral registration in the time domain. Magn Reson Med. 2015 Jan;73(1):44–50. doi: 10.1002/mrm.25094. Epub 2014 Jan 16. PMID: 24436292; PMCID: PMC5851009.

29. Mlynárik V, Cudalbu C, Xin L, Gruetter R. 1H NMR spectroscopy of rat brain in vivo at 14.1Tesla: improvements in quantification of the neurochemical profile. J Magn Reson. 2008 Oct;194(2):163–8. doi: 10.1016/j.jmr.2008.06.019. Epub 2008 Jun 28. PMID: 18703364.

30. Emir UE, Raatz S, McPherson S, Hodges JS, Torkelson C, Tawfik P, White T, Terpstra M. Noninvasive quantification of ascorbate and glutathione concentration in the elderly human brain. NMR Biomed. 2011 Aug;24(7):888–94. doi: 10.1002/nbm.1646. Epub 2011 Jan 12. PMID: 21834011; PMCID: PMC3118919.

31. Mlynárik V, Cudalbu C, Xin L, Gruetter R. 1H NMR spectroscopy of rat brain in vivo at 14.1Tesla: improvements in quantification of the neurochemical profile. J Magn Reson. 2008 Oct;194(2):163–8. doi: 10.1016/j.jmr.2008.06.019. Epub 2008 Jun 28. PMID: 18703364.

32. Gruetter R, Weisdorf SA, Rajanayagan V, Terpstra M, Merkle H, Truwit CL, Garwood M, Nyberg SL, Uğurbil K. Resolution improvements in in vivo 1H NMR spectra with increased magnetic field strength. J Magn Reson. 1998 Nov;135(1):260–4. doi: 10.1006/jmre.1998.1542. PMID: 9799704.

33. Gomez JL, Bonaventura J, Lesniak W, Mathews WB, Sysa-Shah P, Rodriguez LA, Ellis RJ, Richie CT, Harvey BK, Dannals RF, Pomper MG, Bonci A, Michaelides M. Chemogenetics revealed: DREADD occupancy and activation via converted clozapine. Science. 2017 Aug 4;357(6350):503–507. doi: 10.1126/science.aan2475. PMID: 28774929; PMCID: PMC7309169.

34. Manvich DF, Webster KA, Foster SL, Farrell MS, Ritchie JC, Porter JH, Weinshenker D. The DREADD agonist clozapine N-oxide (CNO) is reverse-metabolized to clozapine and produces clozapine-like interoceptive stimulus effects in rats and mice. Sci Rep. 2018 Mar 1;8(1):3840. doi: 10.1038/s41598-018-22116-z. PMID: 29497149; PMCID: PMC5832819

35. Seuwen A, Schroeter A, Grandjean J, Rudin M. Metabolic changes assessed by MRS accurately reflect brain function during drug-induced epilepsy in mice in contrast to fMRI-based hemodynamic readouts. Neuroimage. 2015 Oct 15;120:55–63. doi: 10.1016/j.neuroimage.2015.07.004. Epub 2015 Jul 10. PMID: 26166624.

36. Hughes CL, Stieger KC, Chen K, Vazquez AL, Kozai TDY. Spatiotemporal properties of cortical excitatory and inhibitory neuron activation by sustained and bursting electrical microstimulation. iScience. 2025 May 20;28(6):112707. doi: 10.1016/j.isci.2025.112707. PMID: 40520112; PMCID: PMC12167498.

37. Pasanta D, He JL, Ford T, Oeltzschner G, Lythgoe DJ, Puts NA. Functional MRS studies of GABA and glutamate/Glx - A systematic review and meta-analysis. Neurosci Biobehav Rev. 2023 Jan;144:104940. doi: 10.1016/j.neubiorev.2022.104940. Epub 2022 Nov 2. PMID: 36332780; PMCID: PMC9846867.

38. Sonnay S, Poirot J, Just N, Clerc AC, Gruetter R, Rainer G, Duarte JMN. Astrocytic and neuronal oxidative metabolism are coupled to the rate of glutamate-glutamine cycle in the tree shrew visual cortex. Glia. 2018 Mar;66(3):477–491. doi: 10.1002/glia.23259. Epub 2017 Nov 9. PMID: 29120073.

39. Zhang KJ, Monteggia LM, Kavalali ET. Impact of distinct neurotransmitter release modes on neuronal signaling. Mol Psychiatry. 2026 Feb;31(2):1095–1110. doi: 10.1038/s41380-025-03373-7. Epub 2025 Nov 26. PMID: 41299061; PMCID: PMC12815690

40. McConnell HL, Mishra A. Cells of the Blood-Brain Barrier: An Overview of the Neurovascular Unit in Health and Disease. Methods Mol Biol. 2022;2492:3–24. doi: 10.1007/978-1-0716-2289-6_1. PMID: 35733036; PMCID: PMC9987262.

41. Brookshier A, Lyden P. Differential vulnerability among cell types in the neurovascular unit: Description and mechanisms. J Cereb Blood Flow Metab. 2025 Jan;45(1):3–12. doi: 10.1177/0271678X241299960. Epub 2024 Nov 9. PMID: 39520113; PMCID: PMC11563522.

42. Hall CN, Reynell C, Gesslein B, Hamilton NB, Mishra A, Sutherland BA, O’Farrell FM, Buchan AM, Lauritzen M, Attwell D. Capillary pericytes regulate cerebral blood flow in health and disease. Nature. 2014 Apr 3;508(7494):55–60. doi: 10.1038/nature13165. Epub 2014 Mar 26. PMID: 24670647; PMCID: PMC3976267.

43. Andersen JV. The Glutamate/GABA-Glutamine Cycle: Insights, Updates, and Advances. J Neurochem. 2025 Mar;169(3):e70029. doi: 10.1111/jnc.70029. PMID: 40066661; PMCID: PMC11894596.

44. Pereira M, Vidal B, Valdebenito M, Cheataini F, Bouillot C, Vidal L, Nestor L, De Bundel D, Smolders I, Droguerre M, Zimmer L. Dissociation Between Hemodynamic and Metabolic Responses to Chemogenetic Modulation of Astrocytes in Mouse Visual Cortex. Glia. 2026 Jul;74(7):e70171. doi: 10.1002/glia.70171. PMID: 42206663.

45. Schridde U, Khubchandani M, Motelow JE, Sanganahalli BG, Hyder F, Blumenfeld H. Negative BOLD with large increases in neuronal activity. Cereb Cortex. 2008 Aug;18(8):1814–27. doi: 10.1093/cercor/bhm208. Epub 2007 Dec 5. PMID: 18063563; PMCID: PMC2790390.

46. Arboit A, Krautwald K, Angenstein F. Hemodynamic responses in the rat hippocampus are simultaneously controlled by at least two independently acting neurovascular coupling mechanisms. J Cereb Blood Flow Metab. 2024 Jun;44(6):896–910. doi: 10.1177/0271678X231221039. Epub 2023 Dec 12. PMID: 38087890; PMCID: PMC11318394.

47. Desmond NL, Colbert CM, Zhang DX, Levy WB. NMDA receptor antagonists block the induction of long-term depression in the hippocampal dentate gyrus of the anesthetized rat. Brain Res. 1991 Jun 21;552(1):93–8. doi: 10.1016/0006-8993(91)90664-h. PMID: 1833033.

48. Graves SI, Baker DJ. Implicating endothelial cell senescence to dysfunction in the ageing and diseased brain. Basic Clin Pharmacol Toxicol. 2020 Aug;127(2):102–110. doi: 10.1111/bcpt.13403. Epub 2020 Mar 23. PMID: 32162446; PMCID: PMC7384943.

49. Just N, Xin L, Frenkel H, Gruetter R. Characterization of sustained BOLD activation in the rat barrel cortex and neurochemical consequences. Neuroimage. 2013 Jul 1;74:343–51. doi: 10.1016/j.neuroimage.2013.02.042. Epub 2013 Mar 5. PMID: 23473934.

50. Just N, Sonnay S. Investigating the Role of Glutamate and GABA in the Modulation of Transthalamic Activity: A Combined fMRI-fMRS Study. Front Physiol. 2017 Jan 31;8:30. doi: 10.3389/fphys.2017.00030. PMID: 28197105; PMCID: PMC5281558.

51. Ligneul C, Fernandes FF, Shemesh N. High temporal resolution functional magnetic resonance spectroscopy in the mouse upon visual stimulation. Neuroimage. 2021 Jul 1;234:117973. doi: 10.1016/j.neuroimage.2021.117973. Epub 2021 Mar 21. PMID: 33762216.

52. Roth BL. DREADDs for Neuroscientists. Neuron. 2016 Feb 17;89(4):683-94. doi: 10.1016/j.neuron.2016.01.040. PMID: 26889809; PMCID: PMC4759656.

53. Urban DJ, Roth BL. DREADDs (Designer Receptors Exclusively Activated by Designer Drugs). Annu Rev Pharmacol Toxicol, 2015.

54. Turrigiano G. Homeostatic synaptic plasticity: local and global mechanisms for stabilizing neuronal function. Cold Spring Harb Perspect Biol. 2012 Jan 1;4(1):a005736. doi: 10.1101/cshperspect.a005736. PMID: 22086977; PMCID: PMC3249629.

55. Davis GW. Homeostatic control of neural activity: from phenomenology to molecular design. Annu Rev Neurosci. 2006;29:307–23. doi: 10.1146/annurev.neuro.28.061604.135751. PMID: 16776588.

56. Keck T, Keller GB, Jacobsen RI, Eysel UT, Bonhoeffer T, Hübener M. Synaptic scaling and homeostatic plasticity in the mouse visual cortex in vivo. Neuron. 2013 Oct 16;80(2):327–34. doi: 10.1016/j.neuron.2013.08.018. PMID: 24139037.

57. Emir UE, Raatz S, McPherson S, Hodges JS, Torkelson C, Tawfik P, White T, Terpstra M. Noninvasive quantification of ascorbate and glutathione concentration in the elderly human brain. NMR Biomed. 2011 Aug;24(7):888–94. doi: 10.1002/nbm.1646. Epub 2011 Jan 12. PMID: 21834011; PMCID: PMC3118919.

58. Stiernman L, Grill F, McNulty C, Bahrd P, Panes Lundmark V, Axelsson J, Salami A, Rieckmann A. Widespread fMRI BOLD Signal Overactivations during Cognitive Control in Older Adults Are Not Matched by Corresponding Increases in fPET Glucose Metabolism. J Neurosci. 2023 Apr 5;43(14):2527–2536. doi: 10.1523/JNEUROSCI.1331-22.2023. Epub 2023 Mar 3. PMID: 36868855; PMCID: PMC10082451.

59. Hall CN, Reynell C, Gesslein B, Hamilton NB, Mishra A, Sutherland BA, O’Farrell FM, Buchan AM, Lauritzen M, Attwell D. Capillary pericytes regulate cerebral blood flow in health and disease. Nature. 2014 Apr 3;508(7494):55–60. doi: 10.1038/nature13165. Epub 2014 Mar 26. PMID: 24670647; PMCID: PMC3976267.

60. Epp SM, Castrillón G, Yuan B, Andrews-Hanna J, Preibisch C, Riedl V. BOLD signal changes can oppose oxygen metabolism across the human cortex. Nat Neurosci. 2025 Dec 16. doi: 10.1038/s41593-025-02132-9.

61. Sternson SM, Roth BL. Chemogenetic tools to interrogate brain functions. Annu Rev Neurosci. 2014;37:387–407. doi: 10.1146/annurev-neuro-071013-014048. Epub 2014 Jun 16. PMID: 25002280.

62. Kügler S, Kilic E, Bähr M. Human synapsin 1 gene promoter confers highly neuron-specific long-term transgene expression from an adenoviral vector in the adult rat brain depending on the transduced area. Gene Ther. 2003 Feb;10(4):337–47. doi: 10.1038/sj.gt.3301905. PMID: 12595892.

63. Morrison PD, Jauhar S, Young AH. The mechanism of action of clozapine. J Psychopharmacol. 2025 Apr;39(4):297–300. doi: 10.1177/02698811251319458. Epub 2025 Feb 13. PMID: 39945414; PMCID: PMC11967075.

64. Aggarwal A, Negrean A, Chen Y, Iyer R, Reep D, Liu A, Palutla A, Xie ME, MacLennan BJ, Hagihara KM, Kinsey LW, Sun JL, Yao P, Zheng J, Tsang A, Tsegaye G, Zhang Y, Patel RH, Arthur BJ, Hiblot J, Leippe P, Tarnawski M, Marvin JS, Vevea JD, Turaga SC, Tebo AG, Carandini M, Rossi LF, Kleinfeld D, Konnerth A, Svoboda K, Turner GC, Hasseman JP, Podgorski K. Glutamate indicators with increased sensitivity and tailored deactivation rates. Nat Methods. 2026 Feb;23(2):417–425. doi: 10.1038/s41592-025-02965-z. Epub 2025 Dec 23. PMID: 41436654; PMCID: PMC12904790.

