## Supplemental Tables for "Dissociation between hemodynamic and neurochemical responses during chemogenetic modulation of cortical circuits in rats"

| **Metabolite** | **Comparison Group** | **Dir** | **Δ (%)** | **SD** | **p-value** | **Sig** | **Cohen’s d** |
| --- | --- | --- | --- | --- | --- | --- | --- |
| **Asp** | eD-Ipsi vs iD-Contra | ↑ | +19.0 | 13.6 | <0.01 | ** | 1.40 |
|  | eD-Ipsi vs Sham-Contra | ↑ | +45.8 | 18.7 | *** | *** | 2.45 |
|  | eD-Ipsi vs Saline-Contra | ↓ | -9.0 | 18.4 | ns | ns | -0.49 |
|  | eD-Contra vs iD-Ipsi | ↑ | +36.6 | 14.4 | 0.001 | *** | 3.64 |
|  | eD-Contra vs iD-Contra | ↑ | +61.6 | 14.9 | 0.001 | *** | 5.68 |
|  | iD-Ipsi vs Sham-IPSI | ↓ | -24.8 | 8.1 | 0.001 | *** | -3.86 |
| **GABA** | eD-Ipsi vs iD-Contra | ↑ | +25.8 | 8.1 | <0.001 | *** | 3.18 |
|  | eD-Ipsi vs Sham-Contra | ↑ | +25.9 | 4.5 | *** | *** | 5.76 |
|  | eD-Ipsi vs Saline-Contra | ↑ | +10.6 | 5.2 | ns | ns | 2.04 |
|  | eD-Contra vs iD-Ipsi | ↑ | +33.7 | 19.3 | 0.001 | *** | 2.54 |
|  | eD-Contra vs iD-Contra | ↑ | +43.9 | 18.0 | 0.001 | *** | 3.14 |
|  | iD-Ipsi vs Saline-IPSI | ↓ | -8.5 | 24.0 | ns | ns | -0.57 |
| **Glc** | eD-Ipsi vs iD-Contra | ↑ | +39.2 | 8.6 | <0.001 | *** | 4.56 |
|  | eD-Ipsi vs Sham-Contra | ↑ | +13.4 | 7.8 | ns | ns | 1.72 |
|  | eD-Contra vs iD-Ipsi | ↑ | +25.5 | 18.4 | 0.001 | *** | 2.02 |
|  | iD-Ipsi vs Sham-IPSI | ↓ | -25.9 | 7.5 | 0.001 | *** | -4.45 |
|  | iD-Contra vs Sham-Contra | ↓ | -25.8 | 2.4 | 0.001 | *** | -14.3 |
| **Ins** | eD-Ipsi vs iD-Contra | ↓ | -11.7 | 12.2 | ns | ns | 0.96 |
|  | eD-Contra vs iD-Contra | ↑ | +19.2 | 9.4 | 0.05 | * | 2.68 |
|  | iD-Ipsi vs Saline-IPSI | ↑ | +17.8 | 12.4 | 0.05 | * | 2.19 |
|  | Sham-IPSI vs Saline-IPSI | ↑ | +20.0 | 10.9 | 0.05 | * | 2.59 |
| **GPC+PCh** | eD-Ipsi vs iD-Contra | ↑ | +51.5 | 5.4 | <0.001 | *** | 9.54 |
|  | eD-Ipsi vs Sham-Contra | ↑ | +41.8 | 8.3 | *** | *** | 5.03 |
|  | eD-Ipsi vs Saline-Contra | ↑ | +22.2 | 8.6 | ** | ** | 2.58 |
|  | iD-Ipsi vs Sham-IPSI | ↑ | +26.1 | 10.1 | 0.001 | *** | 3.25 |
|  | iD-Ipsi vs Saline-IPSI | ↑ | +40.8 | 23.7 | 0.001 | *** | 2.68 |
|  | Saline-Ipsi vs Saline-Contra | ↓ | -50.2 | 13.2 | 0.001 | *** | -2.14 |
| **Cr+PCr** | eD-Ipsi vs iD-Contra | ↑ | +21.7 | 3.6 | <0.01 | ** | 6.02 |
|  | eD-Contra vs iD-Ipsi | ↑ | +31.3 | 8.1 | 0.001 | *** | 5.60 |
|  | iD-Ipsi vs Sham-IPSI | ↓ | -25.4 | 4.3 | 0.001 | *** | -7.58 |
|  | iD-Contra vs Sham-Contra | ↓ | -10.8 | 1.0 | 0.05 | ns | -13.2 |
| **Glu+Gln** | eD-Ipsi vs iD-Contra | ↑ | +20.0 | 12.0 | <0.01 | ** | 1.67 |
|  | eD-Contra vs iD-Ipsi | ↑ | +13.2 | 5.0 | 0.05 | ns | 3.78 |
|  | iD-Contra vs Sham-Contra | ↓ | -23.5 | 3.1 | 0.01 | ** | -9.5 |

**Supplementary Table 1: Consolidated Late-Point Metabolic Comparisons (n = 44 comparisons)**

***Note: p-values are Bonferroni-corrected for 44 comparisons. Direction (Dir) and percentage change (Δ %) refer to the first group relative to the second. eDREADDs (eD), iDREADDS (iD)***

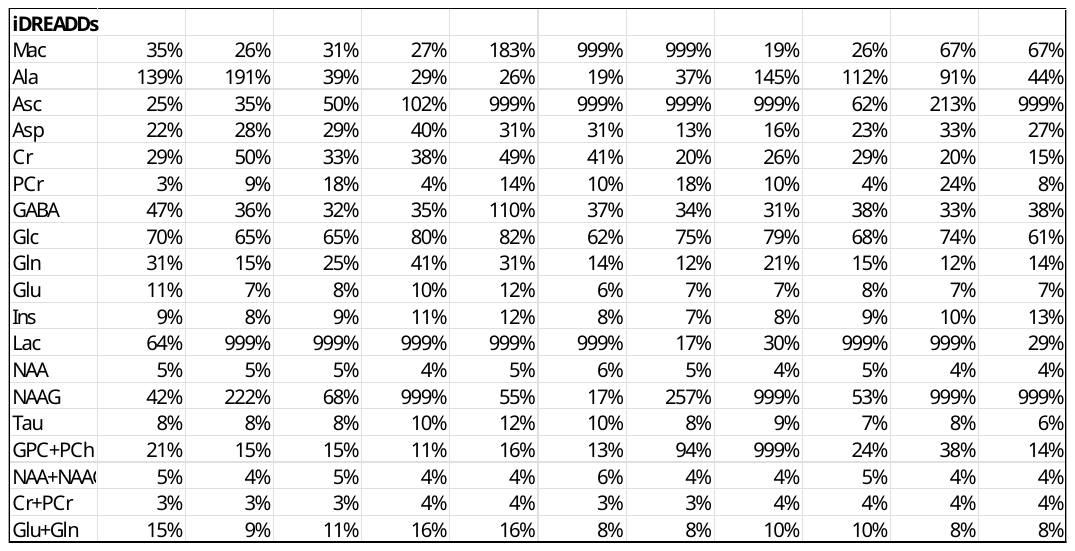

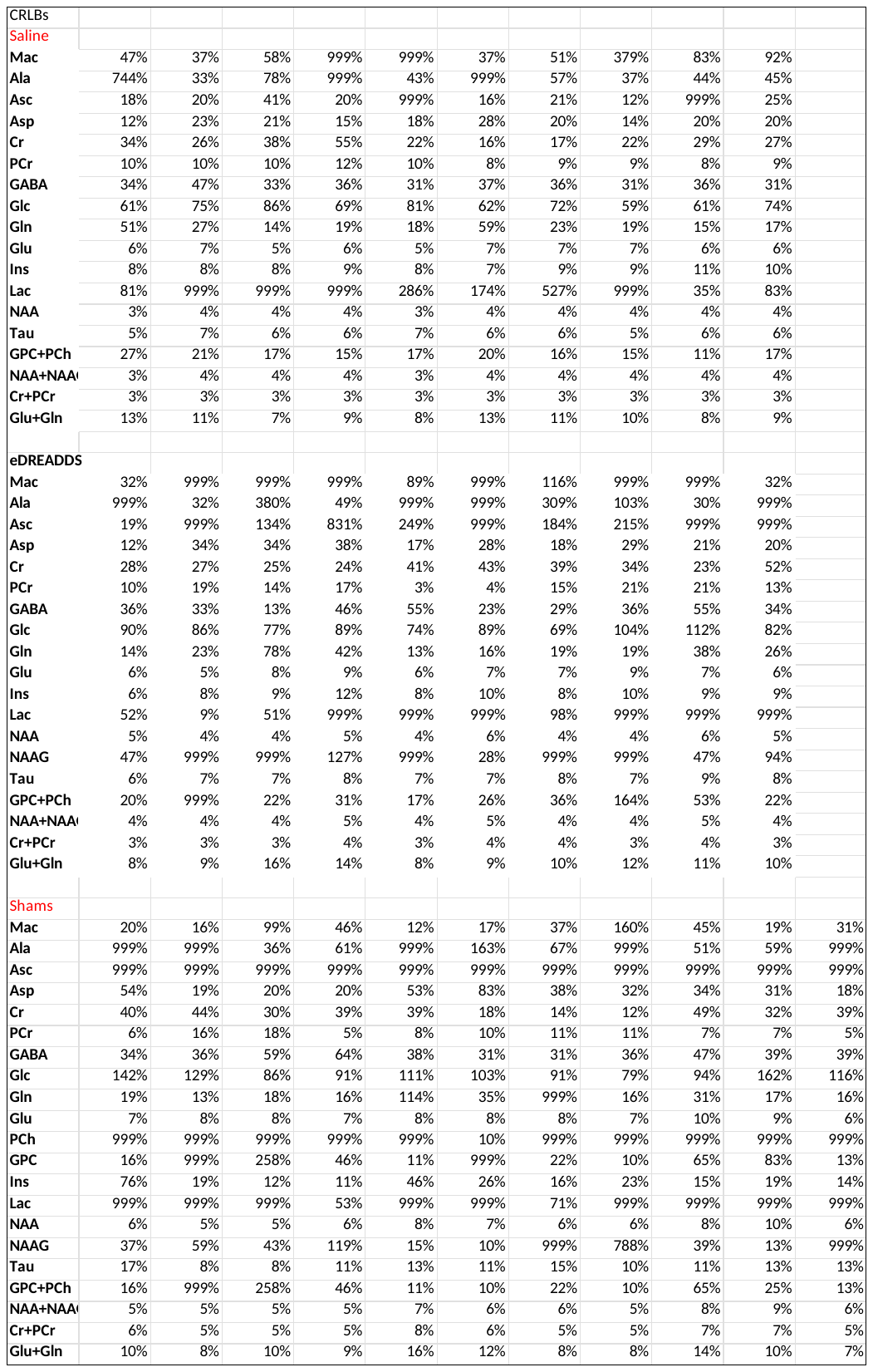
**Mean CRLBs (± Standard Deviations) during functional acquisitions**

**Mean CRLBs ((± Standard Deviations) for eDREADDs, Shams and iDREADDs at 70 min post-CNO at baseline and in the ipsilateral and contralateral brain**

|  | Baseline | SD | IPSI | SD | CONTRA | SD |
| --- | --- | --- | --- | --- | --- | --- |
| Mac | 18% | 5.5 | 52% | 5.2 | 63% | 3.9 |
| Ala | 67% | 0.1 | 84% | 0.4 | 367% | 5.5 |
| Asc | 100% | 0.1 | 275% | 5.7 | 56% | 0.7 |
| Asp | 18% | 0.0 | 20% | 0.1 | 19% | 0.1 |
| Cr | 34% | 0.1 | 21% | 0.0 | 24% | 0.2 |
| PCr | 9% | 0.0 | 9% | 0.0 | 9% | 0.0 |
| GABA | 31% | 0.1 | 30% | 0.1 | 36% | 0.2 |
| Glc | 65% | 0.0 | 69% | 0.2 | 60% | 0.2 |
| Gln | 76% | 0.1 | 32% | 0.1 | 46% | 0.3 |
| Glu | 7% | 0.0 | 6% | 0.0 | 7% | 0.0 |
| Ins | 8% | 0.0 | 6% | 0.0 | 6% | 0.0 |
| Lac | 322% | 0.5 | 522% | 5.6 | 359% | 5.5 |
| NAA | 4% | 0.0 | 3% | 0.0 | 3% | 0.0 |
| NAAG | 30% | 0.1 | 156% | 1.2 | 66% | 0.4 |
| Tau | 6% | 0.0 | 6% | 0.0 | 6% | 0.0 |
| GPC+PCh | 12% | 0.0 | 46% | 0.8 | 15% | 0.1 |
| NAA+NAAG | 4% | 0.0 | 3% | 0.0 | 3% | 0.0 |
| Cr+PCr | 3% | 0.0 | 3% | 0.0 | 3% | 0.0 |
| Glu+Gln | 12% | 0.0 | 11% | 0.0 | 11% | 0.0 |

|  | Baseline | SD | IPSI | SD | CONTRA | SD |
| --- | --- | --- | --- | --- | --- | --- |
| Mac | 62% | 5.48 | 73% | 0.27 | 59% | 5.73 |
| Ala | 75% | 0.49 | 362% | 5.52 | 49% | 0.36 |
| Asc | 351% | 5.61 | 365% | 5.50 | 20% | 0.04 |
| Asp | 19% | 0.04 | 20% | 0.02 | 16% | 0.01 |
| Cr | 27% | 0.13 | 33% | 0.16 | 24% | 0.02 |
| PCr | 9% | 0.03 | 12% | 0.06 | 10% | 0.00 |
| GABA | 32% | 0.17 | 32% | 0.12 | 40% | 0.62 |
| Glc | 63% | 0.10 | 71% | 0.11 | 70% | 0.18 |
| Gln | 27% | 0.20 | 42% | 0.25 | 27% | 0.16 |
| Glu | 6% | 0.00 | 6% | 0.00 | 7% | 0.03 |
| Ins | 7% | 0.03 | 10% | 0.06 | 7% | 0.01 |
| Lac | 73% | 0.34 | 63% | 0.31 | 522% | 6.75 |
| NAA | 3% | 0.01 | 4% | 0.01 | 4% | 0.01 |
| NAAG | 40% | 0.20 | 351% | 5.61 | 32% | 0.08 |
| Tau | 7% | 0.03 | 8% | 0.04 | 8% | 0.02 |
| GPC+PCh | 12% | 0.02 | 14% | 0.08 | 14% | 0.05 |
| NAA+NAAG | 3% | 0.01 | 4% | 0.01 | 4% | 0.01 |
| Cr+PCr | 3% | 0.01 | 3% | 0.01 | 3% | 0.00 |
| Glu+Gln | 9% | 0.02 | 10% | 0.02 | 11% | 0.06 |

**eDREADDs	Shams**

|  | Baseline | SD | IPSI | SD | CONTRA | SD |
| --- | --- | --- | --- | --- | --- | --- |
| Mac | 57% | 4.16 | 54% | 4.9 | 71% | 5.4 |
| Ala | 55% | 0.37 | 69% | 0.5 | 689% | 5.4 |
| Asc | 30% | 0.19 | 206% | 2.5 | 364% | 5.5 |
| Asp | 19% | 0.02 | 20% | 0.0 | 19% | 0.0 |
| Cr | 36% | 0.25 | 23% | 0.0 | 50% | 0.3 |
| PCr | 9% | 0.02 | 7% | 0.0 | 11% | 0.0 |
| GABA | 33% | 0.13 | 36% | 0.0 | 35% | 0.2 |
| Glc | 66% | 0.07 | 62% | 0.1 | 68% | 0.1 |
| Gln | 28% | 0.10 | 26% | 0.1 | 23% | 0.1 |
| Glu | 7% | 0.01 | 8% | 0.0 | 7% | 0.0 |
| GPC | 496% | 4.94 | 999% | 0.0 | 352% | 5.6 |
| Ins | 8% | 0.02 | 9% | 0.0 | 9% | 0.0 |
| NAA | 4% | 0.01 | 4% | 0.0 | 4% | 0.0 |
| NAAG | 108% | 0.98 | 31% | 0.0 | 108% | 0.6 |
| PE | 32% | 0.20 | 30% | 0.0 | 20% | 0.1 |
| Tau | 6% | 0.01 | 7% | 0.0 | 8% | 0.0 |
| GPC+PCh | 11% | 0.02 | 16% | 0.0 | 23% | 0.1 |
| NAA+NAAG | 4% | 0.01 | 4% | 0.0 | 4% | 0.0 |
| Cr+PCr | 3% | 0.01 | 3% | 0.0 | 3% | 0.0 |
| Glu+Gln | 10% | 0.01 | 12% | 0.0 | 10% | 0.0 |

**iDREADDs**
