## Supplemental Figures for "Dissociation between hemodynamic and neurochemical responses during chemogenetic modulation of cortical circuits in rats"

**Supplementary Figure 1**: Mean concentrations of tNAA, Glx, and GABA (mM) and SNR values measured at baseline, transition, and active phases in eDREADDs, iDREADDs, sham, and control groups. Bars represent group means with error bars indicating variability. For tNAA concentrations, plain lines represent trends across successive conditions. Although tNAA decreased in eDREADDs, this trend was not significant.

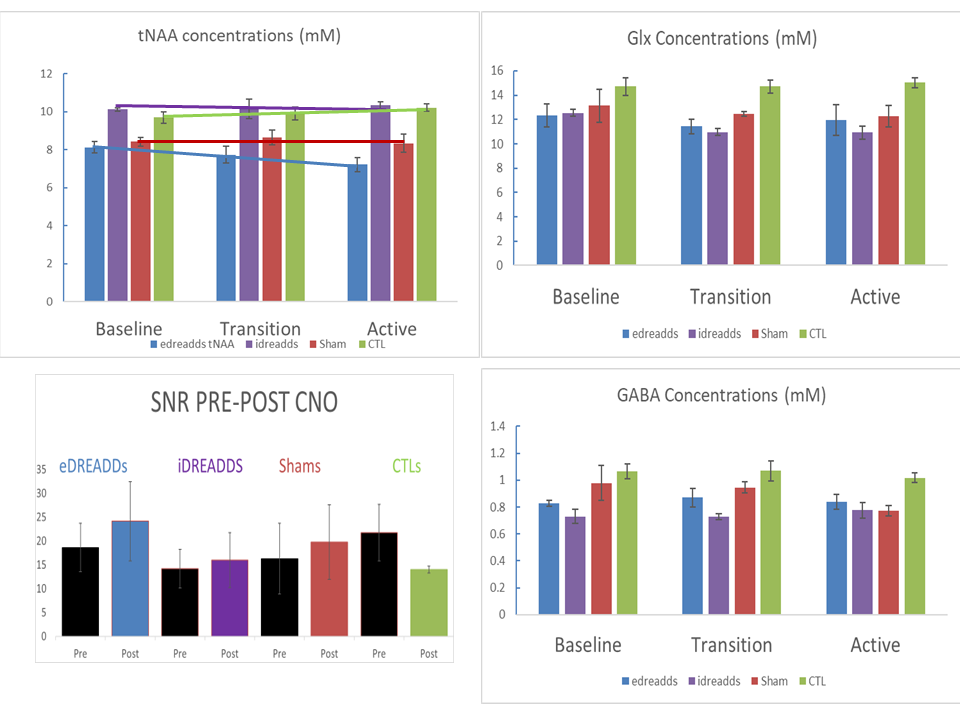

**Suppl.Figure 2. Effect size analysis (Cohen’s *d*) of Glx and GABA time courses highlights dominant baseline group differences**

(A) Time‑resolved Cohen’s *d* values for Glx (glutamate + glutamine) comparisons across the fMRS acquisition period. Effect sizes are shown for eDREADD vs iDREADD (red) and iDREADD vs Sham (green) at successive time windows following CNO administration. Large and persistent effect sizes (|*d*| ≈ 2) are observed for eDREADD vs iDREADD across the entire time course, including baseline, indicating strong chronic metabolic differences between excitatory and inhibitory DREADD groups. In contrast, effect sizes for iDREADD vs Sham remain small to moderate and do not show systematic amplification following CNO injection.(B) Corresponding Cohen’s *d* time courses for GABA show generally smaller effect sizes than for Glx. The largest effects are observed for comparisons involving iDREADD animals at baseline, whereas effect sizes during the CNO challenge remain near zero or fluctuate without a consistent temporal trend, indicating the absence of a robust acute chemogenetic effect on GABA concentrations.Across both metabolites, effect sizes do not increase following CNO administration, demonstrating that the main sources of variance in Glx and GABA concentrations reflect baseline (chronic) group differences rather than acute functional changes. These findings are consistent with the linear mixed‑effects analyses reported in Table 2 and support the interpretation that robust chemogenetically induced BOLD responses are not accompanied by detectable acute shifts in bulk neurometabolite pools at 7 T.
For all groups, *n* =6- 8 rats.

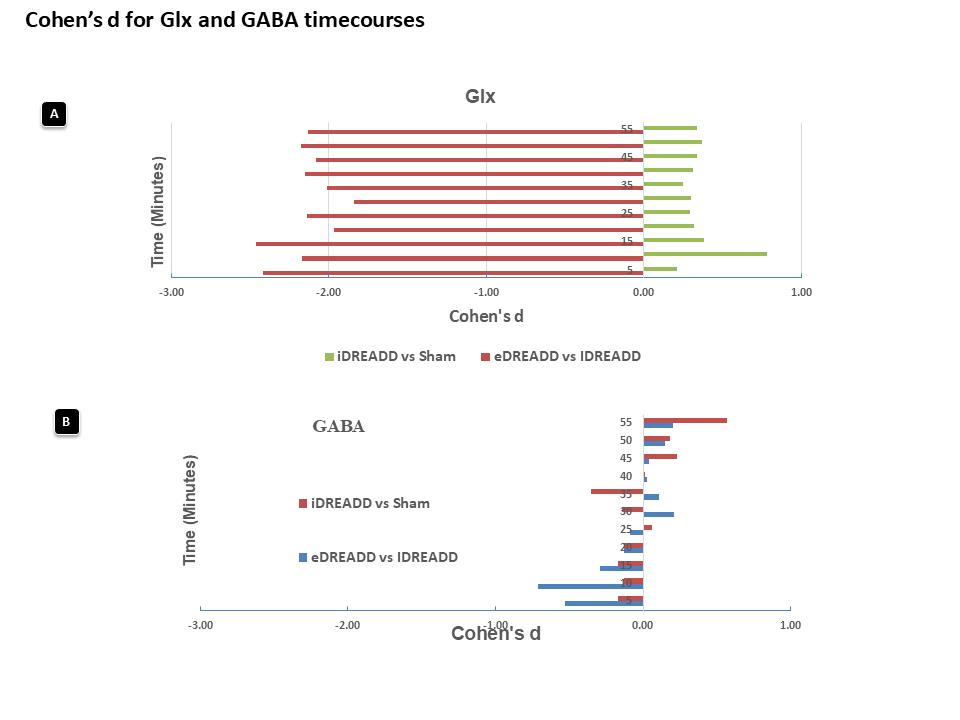

**Suppl. Figure 3. Simulated fMRS time courses demonstrate field‑strength–dependent detectability of chemogenetic metabolic effects**

Simulated ¹H‑fMRS metabolite time courses for Glx (or Glu + Gln) and GABA under identical underlying biological perturbations, shown for 7 T (top) and 14 T (bottom) conditions during excitatory chemogenetic modulation. At 7 T, simulated metabolite signals exhibit substantial temporal variability and almost no visible effect, reflecting reduced signal‑to‑noise ratio. In contrast, simulations at 14 T show improved spectral stability, reduced variance, and clearer separation of individual metabolite components. Right panels depict the corresponding Glx and GABA and time‑resolved Cohen’s *d*, illustrating limited and unstable effect‑size detectability at 7 T compared with consistently larger and more stable effects at 14 T although GABA responses and effect sizes were also limited at 14T. **(B)** Equivalent simulations for inhibitory DREADD–like perturbations further illustrate this field‑strength dependence. At 7 T, Glx/GABA ratio estimates fluctuate substantially and corresponding Cohen’s *d* values remain small and inconsistent despite the presence of an imposed biological effect. At 14 T, the same perturbation yields smoother metabolite trajectories and coherent Glx/GABA ratio changes with larger absolute Cohen’s *d* values, indicating improved sensitivity to excitatory–inhibitory balance shifts. Across both conditions, simulations impose identical underlying neurometabolic changes at 7 T and 14 T, differing only in modelled spectral resolution and noise characteristics. The results indicate that the absence of detectable acute metabolic effects in experimental 7 T fMRS data can be explained by sensitivity limitations rather than by the absence of biological modulation, and predict that similar chemogenetic effects would be detectable at higher magnetic field strengths.**(C-D).** The corresponding SNR versus effect size at 7T and 14T were also depicted. The blue circles identify the SNR region corresponding to this current study.

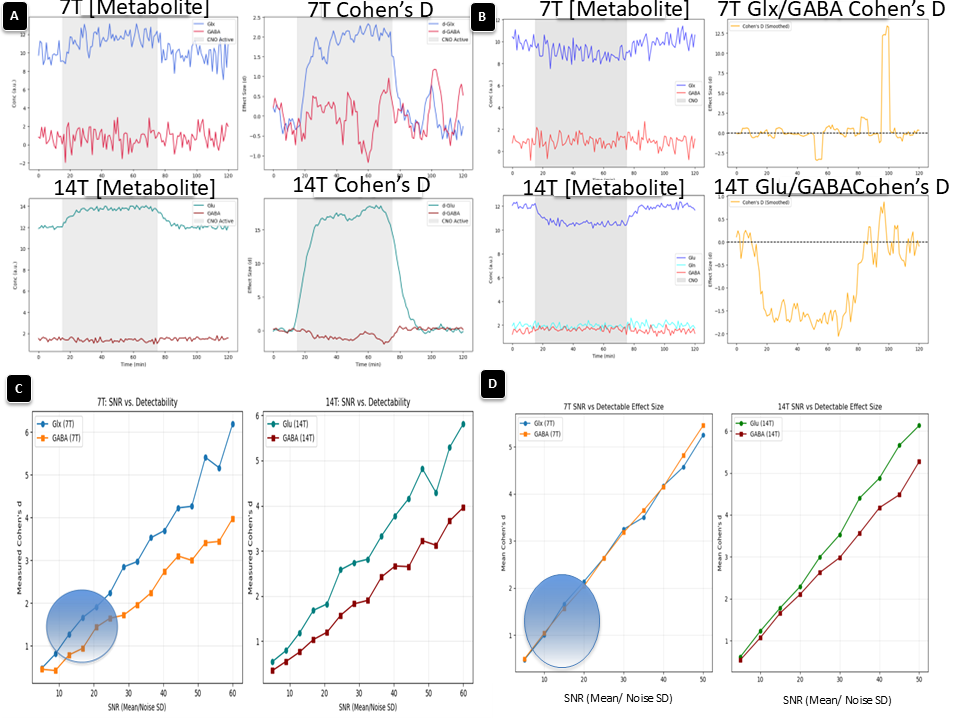
