## Supplementary material for "Dissociation between hemodynamic and neurochemical responses during chemogenetic modulation of cortical circuits in rats": Simulations

**Simulated Detectability of Chemogenetic Metabolic Effects at 7T and 14T**

At 7T, simulated Glx and GABA concentration time courses exhibited substantial variability and low SNR with limited changes during excitatory (Suppl. Fig.3A) or inhibitory modulations (Suppl. Fig 3B). Despite the presence of an underlying chemogenetic modulation in the model, effect sizes remained small with time-resolved Cohen’s d values largely below thresholds typically considered biologically meaningful except for Glx at 7T (-2 to 2). The Cohen’s d values versus SNR were also simulated at 7T and 14T for both the excitatory and inhibitory cases with the range of SNR values found in the present study (Fig 3, blue Circle). These results closely mirror the experimental ^1^H-fMRS findings obtained at 7T, where no robust metabolic changes were observed during the expected DREADD activation window. In contrast, simulations at 14T revealed markedly improved detectability of metabolic changes for Glu but not for GABA. Increased signal-to-noise ratio and improved spectral separation allowed Glu and GABA-related signals to exhibit smoother time courses and reduced variance across simulated subjects. Under identical biological perturbations, Glu/GABA ratios at 14T showed consistent deviations from baseline, with effect sizes exceeding those observed at 7T (< -2 or > 2). These findings suggest that chemogenetic modulation of excitatory–inhibitory balance could be detectable using ^1^H-fMRS at higher magnetic field strengths, even when remaining below detection limits at 7T but remained to be further explored.

Mathematical equations that directly led to the script and the simulation of metabolite time courses during CNO activation of DREADDs. This script simulates metabolite dynamics (Glx, Glu, Gln, GABA) under inhibitory DREADD activation using first-order differential Equations with simple linear dynamics toward a modulated target.

**General Concept**

For each metabolite $M$, the concentration over time is modeled as:

$$\frac{dM}{dt}=\frac{M_{\text{target}}(t)-M(t)}{\tau_{M}}$$

Where:

- $M_{\text{target}}(t)=M_{0}\cdot[1+\text{fractional change due to CNO}(t)]$
- $\tau_{M}$= time constant (controls speed of adaptation toward the target)
- $M_{0}$= baseline metabolite concentration
- Fractional change depends on **CNO activation**, which is on/off depending on time.

**CNO Activation Function**

The CNO drug is modeled as a **binary switch**:

$$\text{CNO}(t)=\left\{ \begin{matrix} 1 & \text{if }CNO_{\text{onset}}\leq t\leq CNO_{\text{onset}}+CNO_{\text{duration}} \\ 0 & \text{otherwise} \end{matrix} \right.$$

Fractional change is:

$$\text{frac}_{M}(t)=E_{\max,M}\cdot\text{CNO}(t)$$

Where $E_{\max,M}$is the maximal effect (positive or negative).

**7T Differential Equations**

Two metabolites: Glx and GABA.

$$\begin{matrix} \frac{d\text{Glx}}{dt} & =\frac{\text{Glx}_{0}\cdot[1+E_{\max,\text{Glx}}\cdot\text{CNO}(t)]-\text{Glx}}{\tau_{\text{Glx}}} \\ \frac{d\text{GABA}}{dt} & =\frac{\text{GABA}_{0}\cdot[1+E_{\max,\text{GABA}}\cdot\text{CNO}(t)]-\text{GABA}}{\tau_{\text{GABA}}} \end{matrix}$$

- Here, $E_{\max,\text{Glx}}=-0.1$(Glx decreases), $E_{\max,\text{GABA}}=0.1$(GABA increases).

**14T Differential Equations**

Three metabolites: Glu, Gln, GABA.

$$\begin{matrix} \frac{d\text{Glu}}{dt} & =\frac{\text{Glu}_{0}\cdot[1+E_{\max,\text{Glu}}\cdot\text{CNO}(t)]-\text{Glu}}{\tau_{\text{Glu}}} \\ \frac{d\text{Gln}}{dt} & =\frac{\text{Gln}_{0}\cdot[1+E_{\max,\text{Gln}}\cdot\text{CNO}(t)]-\text{Gln}}{\tau_{\text{Gln}}} \\ \frac{d\text{GABA}}{dt} & =\frac{\text{GABA}_{0}\cdot[1+E_{\max,\text{GABA}}\cdot\text{CNO}(t)]-\text{GABA}}{\tau_{\text{GABA}}} \end{matrix}$$

- Parameter values:
  - $E_{\max,\text{Glu}}=-0.12$
  - $E_{\max,\text{Gln}}=-0.03$
  - $E_{\max,\text{GABA}}=0.1$

**Summary of Dynamics**

- Each metabolite **relaxes exponentially** toward a new target set by the baseline × (1 + fractional effect).
- Time constant $\tau$sets **how fast** this relaxation occurs.
- The **CNO binary function** determines when the perturbation happens.
- Noise is added afterward for realism: $M_{\text{observed}}=M_{\text{ODE}}+\text{Gaussian noise}$.

**Optional: Compact Form**

For a metabolite $M$:

$$\frac{dM}{dt}=\frac{M_{0}(1+E_{\max,M}\cdot\mathbf{1}_{\left[ t_{\text{onset}},t_{\text{onset}}+t_{\text{duration}} \right]}(t))-M}{\tau_{M}}$$

Where $\mathbf{1}_{\left[ a , b \right]}(t)$is the **indicator function** (1 if $t\in[a,b]$, else 0).
